# Multi-modal locomotor costs explain sexual size but not shape dimorphism in a leaf-mimicking insect

**DOI:** 10.1101/2021.07.06.451325

**Authors:** Romain P. Boisseau, Thies H. Büscher, Lexi J. Klawitter, Stanislav N. Gorb, Douglas J. Emlen, Bret W. Tobalske

**Author notes:** Corresponding author: RB.

## Abstract

In most arthropods, adult females are larger than males, and male competition is a race to quickly locate and mate with scattered females (scramble competition polygyny). In this context, smaller males may be favored due to more efficient locomotion leading to higher mobility during mate searching while larger males may benefit from increased speed and higher survivorship. Understanding how body size affects different aspects of the locomotor performance of males is therefore essential to shed light on the evolution of this widespread mating system. Using a combination of empirical measures of flight performance and substrate adhesion, and modelling of body aerodynamics, we show that large body size impairs both flight and landing (attachment) performance in male leaf insects (*Phyllium philippinicum*), a species where relatively small and skinny males fly through the canopy in search of large sedentary females. Smaller males were more agile in the air, ascended more rapidly during flight, and had a lower risk of detaching from the substrates on which they walk and land. Our models revealed variation in body shape affected body lift and drag, but tradeoffs with weight meant that effects were negligible, suggesting that flight costs do not explain the evolution of strong sexual dimorphism in body shape in this species.

**Graphical abstract:** 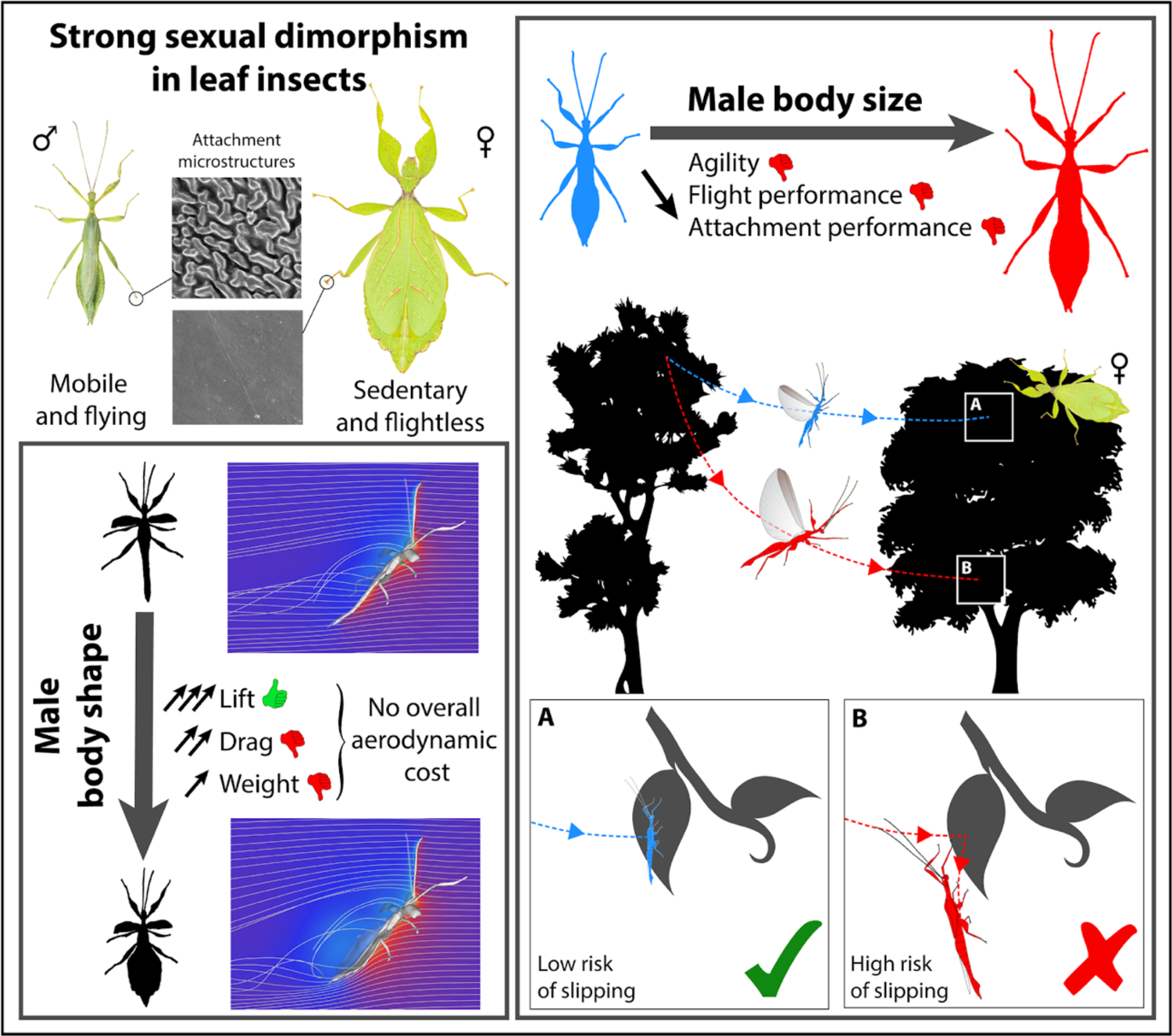

## Introduction

Sexual dimorphism is the ultimate result of sex-dependent selection leading traits toward different optima in each sex (Blanckenhorn, 2005; Cox and Calsbeek, 2009; Fairbairn et al., 2007). Many key hypotheses for the ultimate drivers of sexual dimorphism, and specifically sexual size dimorphism (SSD), remain controversial or relatively poorly supported (Fairbairn et al., 2007). While factors favoring larger bodies are widely recognized –i.e., fecundity selection in females (Honěk, 1993; Pincheira-Donoso and Hunt, 2017) and sexual selection in males (Andersson, 1994) – the selective pressures favoring smaller sizes have received less attention (Blanckenhorn, 2005, 2000; Fairbairn et al., 2007; Kingsolver and Pfennig, 2004). In resource or female defense mating systems, the largest, most armored males are often favored against rivals over access to mating (Andersson, 1994; Emlen, 2014; Emlen and Oring, 1977; Hardy and Briffa, 2013; Shuker and Simmons, 2014). However, when females are dispersed throughout the landscape and do not rely on an easily defensible resource, male competition unfolds instead as a race to locate females (i.e., scramble competition polygyny) (Herberstein et al., 2017). In this context, selection is predicted to favor male traits that increase the distance travelled during mate searching (i.e., mobility) (Herberstein et al., 2017).

A small body size is often assumed to be beneficial for the searching performance of scrambling males (Blanckenhorn, 2000; Kelly, 2020; Kelly et al., 2008) – *e.g.,* by increasing endurance and enabling longer searching times, eventually leading to higher encounter rates with females (Husak and Fox, 2008). For instance, in aerial species, heavier bodies, which have a higher wing loading 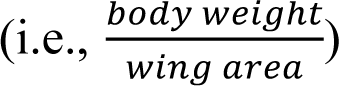 are usually thought to reduce flight performance as they require relatively more power to be maintained in the air, particularly during hovering or slow forward flight (Biewener and Patek, 2018). However, directional selection for smaller sizes does not appear universal among scrambling systems (Herberstein et al., 2017). In some species, it is the larger males that are more mobile and more successful than smaller counterparts, possibly owing to higher survivorship during mate searching, higher movement speeds or larger, more effective sensory structures (Barry, 2013; Hanks et al., 1996). Identifying how different aspects of locomotion are affected by body morphology is therefore critical to understand the variation in the effect of size on mobility. Here, we thoroughly test the hypothesis that smaller males have multiple locomotor advantages during mate searching in a scramble-competition insect species with pronounced sexual dimorphism in size and body shape.

We quantified the effects of variation in morphology (body size and shape) on flight and substrate attachment (landing) performance in male leaf insects (*Phyllium* (*Phyllium*) *philippinicum*, Hennemann, Conle, Gottardo & Bresseel, 2009, Phylliidae, Phasmatodea). In this solitary canopy-dwelling species, large, sedentary adult females are outstanding leaf-mimics due to lateral ‘leaf-like’ expansions of the abdominal segments and legs (Bradler and Buckley, 2018; Hennemann et al., 2009). Adult males are nine times lighter and almost two times slenderer than females, and have relatively longer antennae (i.e., strong size [SSD] and shape dimorphism; Fig. 1). Adult females lack hindwings but have extended forewings that lie flat on their dorsum, aiding in camouflage (Hennemann et al., 2009). These wings cannot flap and can only contribute to parachuting if falling. In contrast, adult males have rudimentary forewings and long, fully-developed transparent hindwings allowing flapping flight. Males use their wings and long antennae to actively search for sedentary females widely scattered in the canopy (Missbach et al., 2014), suggesting that mobility and flight performance may be critical for male fitness (Herberstein et al., 2017).

**Figure 1:**
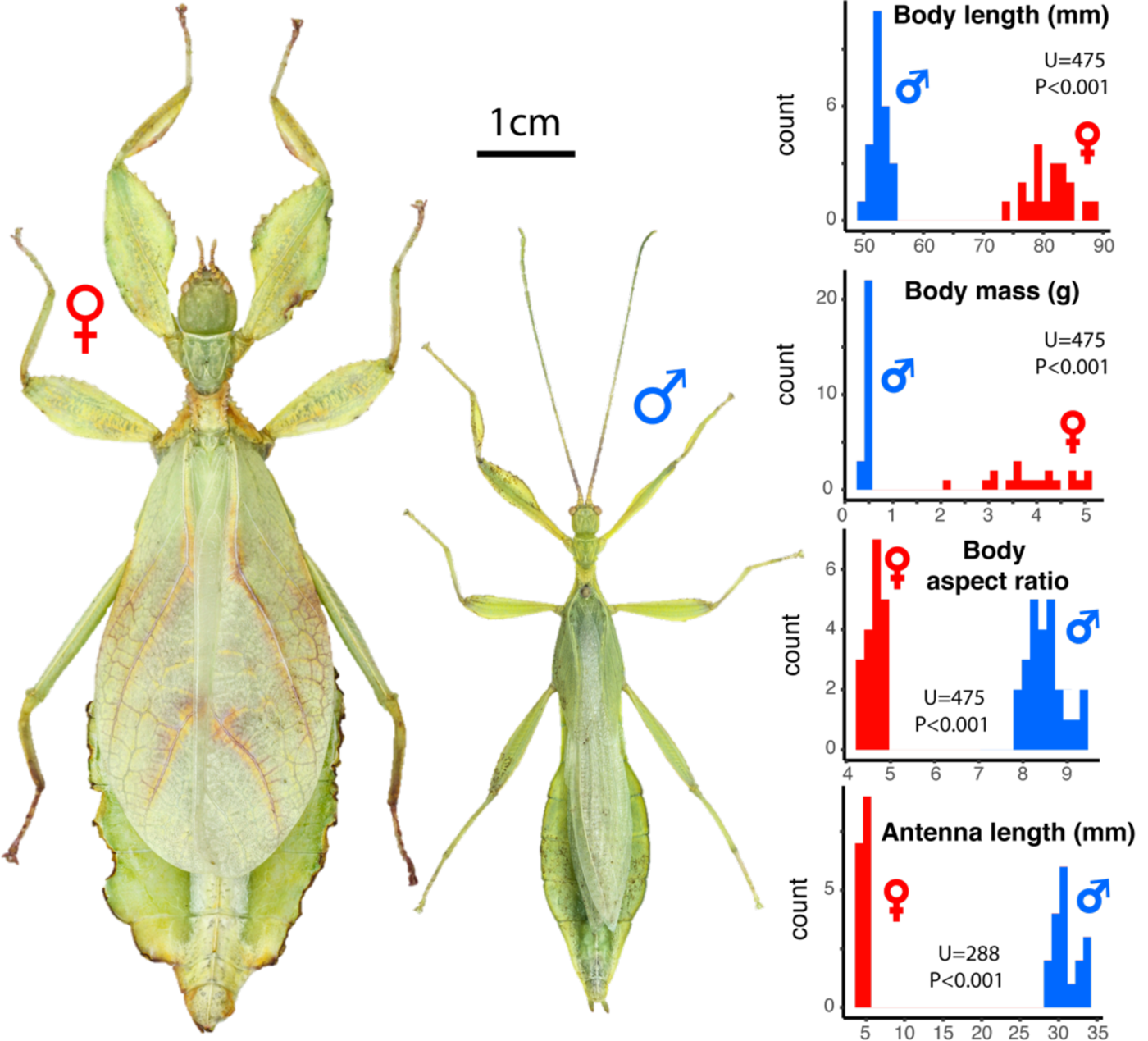
Sexual dimorphism in *P. philippinicum.* Pictures display an adult female (left) and male (right) in dorsal view. Distributions of male (blue) and female (red) body length, mass, aspect ratio and antenna length are shown on the right. Wilcoxon-Mann-Whitney tests are presented to compare the two sexes.

We predicted that increases in body weight would impair flight performance (agility and ascending; Maginnis, 2006; Zeng et al., 2020), and reduce the ability of these insects to hold on to branches or leaves when they landed, increasing the risk of crashing and falling from high perches. We also hypothesized that selection for efficient locomotion (higher lift:drag ratios, lower power requirements) could account for the extensive sexual dimorphism in body shape (the “skinniness” of males relative to females). As these insects fly at low body angles of attack, we predicted that wide ‘leaf-like’ body silhouettes would have proportionally greater drag and, hence, a lower lift to drag ratio, requiring relatively more mechanical work for flying and putting wider males at a locomotor disadvantage. We tested these predictions using an integrative approach combining scaling of gross morphology, micro-structure descriptions, empirical measures of flight and attachment, and modeling of flight costs.

## Results

### Small males were more agile and climbed faster during flight than larger males

Compared to females, male *P. philippinicum* are shorter, lighter, skinnier and have much longer antennae (Fig. 1, S1-2). The large, leaf mimicking females are incapable of flight. For males, wing loading 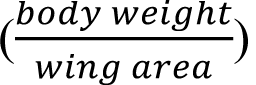 increased significantly with body length (BL) (Fig. S2H, Table S1) but relative flight muscle mass did not (Fig. S2I, Table S1), indicating that large males were not compensating for their relatively heavier weight by building disproportionately large wings or flight muscles. This resulted in a substantial reduction in flight performance, which we assessed using high-speed (500fps) video recordings of flight trajectories (Fig. S3, Video S1-3).

We first measured the rotational velocity of the insect long-axis body angle (i.e., pitch) when recovering from being dropped (see methods, Fig. S4B). The rotation occurred over multiple wingbeats (Video S1-S2), so we interpret it to represent torsional agility, an aspect of aerial maneuverability that reflects how fast the animal can correct its body pitch in the air from a free falling, head first, position to a stable body pitch (Dudley, 2002, 2000). This is distinct from oscillation in body pitch within and among wingbeats which may reflect longitudinal instability and a lack of control (Taylor and Thomas, 2003). We also measured mean horizontal and vertical velocity of the body center of mass to quantify the capacity of the individual to fly forward and ascend. Body mass and wing area had significant opposing effects on rotational velocity and on mean vertical velocity (Table 1). Consistently, wing loading negatively affected rotational velocity (*X*^2^=13.0, df=1, p=0.0003, Fig. 2B) and mean vertical velocity (*X*^2^=11.4, df=1, p=0.0007, Fig. 2C). Thus, lighter males with relatively larger wings --i.e., with a lower wing loading -- were both more adept at changing body pitch angle in the air and had a greater capacity for ascending flight than heavier males with relatively smaller wings (Fig. 2A). Despite flapping their wings at a higher frequency (Table 1, Fig. 2D), larger males also decreased stroke amplitude (Table 1, Fig. 2E) resulting in a comparable wing angular velocity to smaller males that did not permit them to compensate their weight by flying faster (Table 1) and that consequently impaired their agility and ability to climb.

**Table 1:** Analyses of the effects of body mass, wing area and body aspect ratio on various components of flight performance. The table shows a summary of linear mixed effect model outputs including individual ID as a random factor. Fixed effects were mean-centered and standardized. Outputs include the estimated parameter value (± standard error) and a type-I likelihood ratio test to investigate the significance of each fixed effect sequentially. Fixed effects that were found to have a significant effect (p<0.05) are bolded.

| Response variable | Fixed effects | $\beta \pm SE$ | Df | $\chi^2$ | p |
| --- | --- | --- | --- | --- | --- |
| Rotational velocity | **Body mass** | **$-82.7 \pm 25.2$** | **1** | **7.04** | **0.008** |
|  | **Wing area** | **$53.6 \pm 22.9$** | **1** | **5.99** | **0.014** |
| | Body aspect ratio | $13.4 \pm 19.3$ | 1 | 0.63 | 0.43 |
| Mean vertical velocity | **Body mass** | **$-0.37 \pm 0.12$** | **1** | **7.49** | **0.006** |
|  | **Wing area** | **$0.22 \pm 0.10$** | **1** | **4.99** | **0.025** |
| | Body aspect ratio | $0.05 \pm 0.09$ | 1 | 0.40 | 0.53 |
| Mean horizontal velocity | Body mass | $0.23 \pm 0.17$ | 1 | 1.56 | 0.21 |
| | Wing area | $-0.10 \pm 0.15$ | 1 | 0.49 | 0.49 |
| | Body aspect ratio | $0.07 \pm 0.13$ | 1 | 0.44 | 0.51 |
| Wing beat frequency | **Body mass** | **$0.51 \pm 0.29$** | **1** | **4.99** | **0.03** |
| | Wing area | $0.26 \pm 0.26$ | 1 | 1.20 | 0.27 |
|  | **Body aspect ratio** | **$0.43 \pm 0.22$** | **1** | **4.34** | **0.04** |
| Wing beat amplitude | **Body mass** | **$-2.59 \pm 4.37$** | **1** | **4.55** | **0.03** |
| | Wing area | $-6.12 \pm 3.97$ | 1 | 2.93 | 0.09 |
| | Body aspect ratio | $-0.91 \pm 3.36$ | 1 | 0.10 | 0.75 |

**Figure 2:**
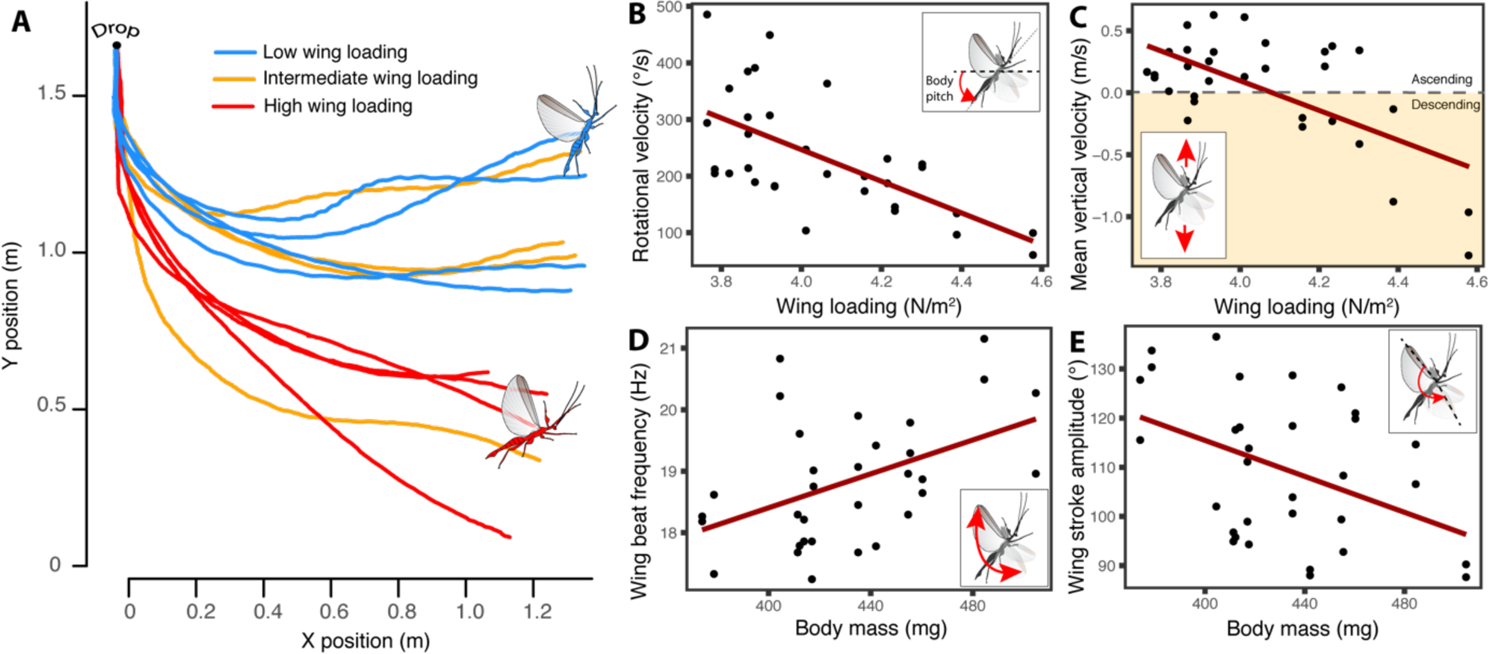
Effect of body mass and wing loading on male *P. philippinicum* flight performance. **A:** Flight trajectories of a subset of males with low (< 4 N.m^-2^), intermediate (4 < WL < 4.22 N.m^-2^) and high (> 4.22 N.m^-2^) wing loading. Rotational velocity (**B**) and mean vertical velocity (**C**) as a function of wing loading. Wingbeat frequency (**D**) and wing stroke amplitude (**E**) as a function of body mass. Mixed effect linear regressions including individual ID as a random factor are fitted in **B, C, D** and **E**.

### Small males attach to substrates better than larger males, reducing their risk of falling

Safely attaching to leaves and branches with their tarsi is critical for canopy-dwelling leaf insects, which are large enough to suffer serious injury from falls. Females spend most of their time hanging from the undersides of leaves and therefore rely on friction attachment forces, which resists tarsal movement along (parallel to) the surface of the leaf, and adhesion forces, which resists falling from the underside of the leaf (i.e., perpendicular to the leaf surface). Male attachment performance also depends on both friction and adhesion forces, but males walk greater distances through the canopy in their search for females, requiring them to attach to a wider variety of plant surfaces (e.g., leaves, branches, trunks), and flying males must overcome inertia and impact forces to hold on to branches or leaves when they land.

Friction (parallel to the substrate) and adhesion (perpendicular) forces are the product of maximum tarsal pad frictional or adhesive strength and pad area. Tarsi of male and female *P. philippinicum* are similar in overall morphology. Both sexes have five-segmented tarsi, each of the tarsomeres equipped with a euplantula (i.e., “heel” pads), and an arolium (i.e., “toe” pads) and two claws on the pretarsus (Fig. 3A), as is typical for phasmids (Büscher et al., 2018b, 2018a). The primary region of tarsal friction is the euplantula, and for adhesion is the arolium (Labonte and Federle, 2013). The overall size of these tarsal pads scales isometrically with BL and does not differ between sexes after accounting for size differences (Table 2, Fig. 3D). However, scanning electron microscopy (SEM) revealed that the attachment microstructures on the euplantulae are sexually dimorphic. The euplantulae of females are smooth, without cuticular microstructures (Fig. 3B), while in males this surface is covered with maze-like arrangements of ridges (Fig. 3C; *sensu* Büscher et al., 2019, 2018b, 2018a). These cuticular microstructures are likely to perform better, on average, on a broad range of substrate surfaces males experience from active searching. In contrast, smooth euplantulae are specifically adapted to smooth surfaces such as the surface of smooth leaves (Büscher and Gorb, 2019; Bußhardt et al., 2012).

**Figure 3:**
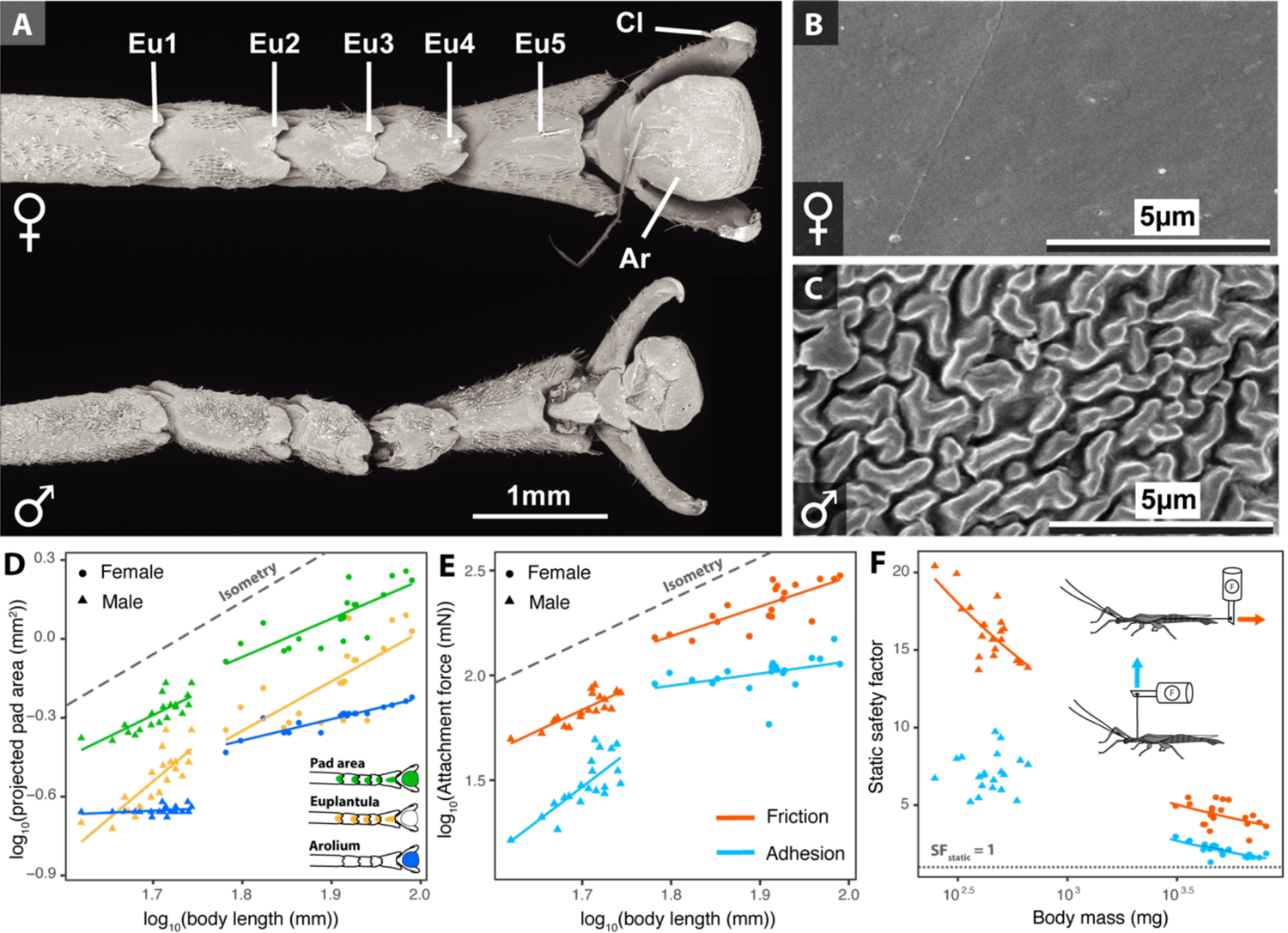
(A) Morphology of female (top) and male (bottom) *P. philippinicum* metathoracic tarsi. Eu. euplantula, Cl, claw, Ar, arolium. Euplantular attachment microstructures are smooth in females (**B**) and bear maze-like cuticular patterns in males (**C**). (**D**) Scaling relationships between body length and adhesive pad areas. (**E**) Friction and adhesion forces as a function of body length. Dashed lines in (**D**) and (**E**) represent the slope expected under isometry (slope = 2, arbitrary intercept). (**F**) Static safety factor as a function of body mass. The x-axis is on a log_10_ scale. The dotted line in (**F**) represent a safety factor of one. Diagrams represent the experimental set-up to measure friction (orange) and adhesion (blue) forces.

**Table 2:** Results of type I ANOVA from linear models contrasting the differences between sexes in terms of scaling relationships between body size and attachment pad areas, attachment forces and static safety factors. Scaling exponents β and the corresponding 95% confidence intervals are shown in comparison to isometric expectations. Significant effects (i.e., p <0.05) are bolded.

| Response variable<br>(log<sub>10</sub> transformed) | Explanatory<br>variables | F | df1 | df2 | P | Isometric<br>slope | Slope β<br>[95% CI] |
| --- | --- | --- | --- | --- | --- | --- | --- |
| **Pad morphology** |  |  |  |  |  |  |  |
| Arolium area | **log<sub>10</sub>(body length)** | **3460** | **1** | **36** | **<0.001** | 2 | Males: 0.15 [-0.07, 0.37]<br>Females: 0.81[0.63, 0.99] |
|  | **sex** | **245.2** | **1** | **36** | **<0.001** |  |  |
|  | **interaction** | **19.0** | **1** | **36** | **<0.001** |  |  |
| Euplantula area | **log<sub>10</sub>(body length)** | **225.6** | **1** | **36** | **<0.001** | 2 | Males: 2.86 [1.87, 3.86]<br>Females: 1.88 [1.00, 2.77] |
|  | sex | 0.42 | 1 | 36 | 0.52 |  |  |
|  | interaction | 1.81 | 1 | 36 | 0.19 |  |  |
| Combined pad area | **log<sub>10</sub>(body length)** | **483.0** | **1** | **36** | **<0.001** | 2 | Males: 1.65 [1.01, 2.28]<br>Females: 1.46 [0.90, 2.02] |
|  | sex | 2.41 | 1 | 36 | 0.13 |  |  |
|  | interaction | 0.17 | 1 | 36 | 0.68 |  |  |
| **Attachment forces** |  |  |  |  |  |  |  |
| Adhesion forces | **log<sub>10</sub>(body length)** | **444.4** | **1** | **36** | **<0.001** | 2 | Males: 3.22 [1.93, 4.51]<br>Females: 0.58[-0.03, 1.18] |
|  | **sex** | **24.3** | **1** | **36** | **<0.001** |  |  |
|  | **interaction** | **16.5** | **1** | **36** | **<0.001** |  |  |
| Friction forces | **log<sub>10</sub>(body length)** | **803.4** | **1** | **36** | **<0.001** | 2 | Males: 1.98 [1.34, 2.63]<br>Females: 1.41 [0.86, 1.97] |
|  | **sex** | **18.3** | **1** | **36** | **<0.001** |  |  |
|  | interaction | 1.54 | 1 | 36 | 0.22 |  |  |
| Static adhesion<br>safety factor | **log<sub>10</sub>(body mass)** | **562.2** | **1** | **36** | **<0.001** | 0 | Males: 0.06 [-0.3, 0.43]<br>Females: -0.58 [-0.85, -0.31] |
|  | sex | 3.5 | 1 | 36 | 0.068 |  |  |
|  | **interaction** | **9.11** | **1** | **36** | **0.005** |  |  |
| Static friction<br>safety factor | **log<sub>10</sub>(body mass)** | **980.9** | **1** | **36** | **<0.001** | 0 | Males: -0.34 [-0.51, -0.17]<br>Females: -0.33 [-0.64, -0.02] |
|  | **sex** | **5.31** | **1** | **36** | **0.027** |  |  |
|  | interaction | 0.008 | 1 | 36 | 0.93 |  |  |

To test for effects of body size on attachment performance, we measured maximal tarsal friction and adhesion using a force transducer mounted on a motorized micromanipulator. Static safety factors (SF_static_) were calculated from measures of the force required to pull the insect horizontally off a glass plate (friction forces), or backwards off the plate (adhesion forces), divided by weight (see methods). This approximates how many times leaf insects can attach their own weight to smooth canopy surfaces when they are resting or hanging motionless, which is what females do most of the time. Flying males, in contrast, experience additional attachment demands when they land, due to impact forces. Consequently, we estimated the typical impact forces experienced by males (Fi) when landing on a leaf based on leaf deflection and landing speed (see methods) to compute dynamic safety factors 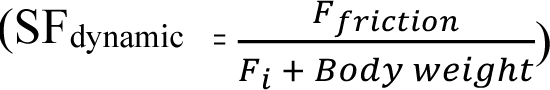 Only frictional attachment forces (as opposed to adhesive forces) were used to compute these SF_dynamic_ as they are the most important for accommodating deceleration and impact forces when landing (Higham et al., 2017).

Larger males displayed higher adhesion and friction attachment forces than smaller ones. This positive correlation between attachment force and BL was also found in females, but only when considering friction forces (Fig. 3E, Table 2). Females also had relatively higher adhesion and friction attachment forces than males (Fig. 3E, Table 2). However, females, with their large, leaf-mimicking bodies and mainly motionless behavior, had the lowest static safety factors (Fig. 3F, Table 2). As they were much lighter, males had relatively larger friction SF_static_ than females (Fig. 3F, Table 2), consistent with the greater attachment demands they experience from active searching and, especially, landing. While males displayed relatively higher adhesion SF_static_, this difference was not significant (p = 0.068, Fig. 3F, Table 2). In females, adhesion and friction SF_static_ negatively correlated with body mass, while in males, only friction SF_static_ significantly decreased with increasing body mass (Fig. 3F, Table 2).

In our landing simulations (see methods), heavier males caused larger leaf deflections when landing as estimated by equation 3 (Fig. 4A,B). Leaf deflection was greatest for the largest leaves (Fig. 4A,B). We found, in our flight experiments, that overall landing velocity was positively correlated with male body mass (Wald chisquare test: *X*^2^ = 4.12, df = 1, p = 0.042). We then used the fixed effect estimates of the corresponding linear mixed model to predict the landing velocity of males given their body mass and estimate impact forces at landing (Fig 4A). Landing impact forces increased with male body mass and were relatively higher for smaller leaves (Fig. 4C, Table 3). Finally, SF_dynamic_ decreased with male body mass and were relatively lower for smaller leaves (Fig. 4D, Table 3). Interestingly, SF_dynamic_ fell below 1.0 for body masses > 350 mg and small leaves (Fig 4D), therefore predicting slippage. In other words, large males are more likely to slip and fall when landing on canopy substrates, especially on smaller and stiffer leaves.

**Figure 4:**
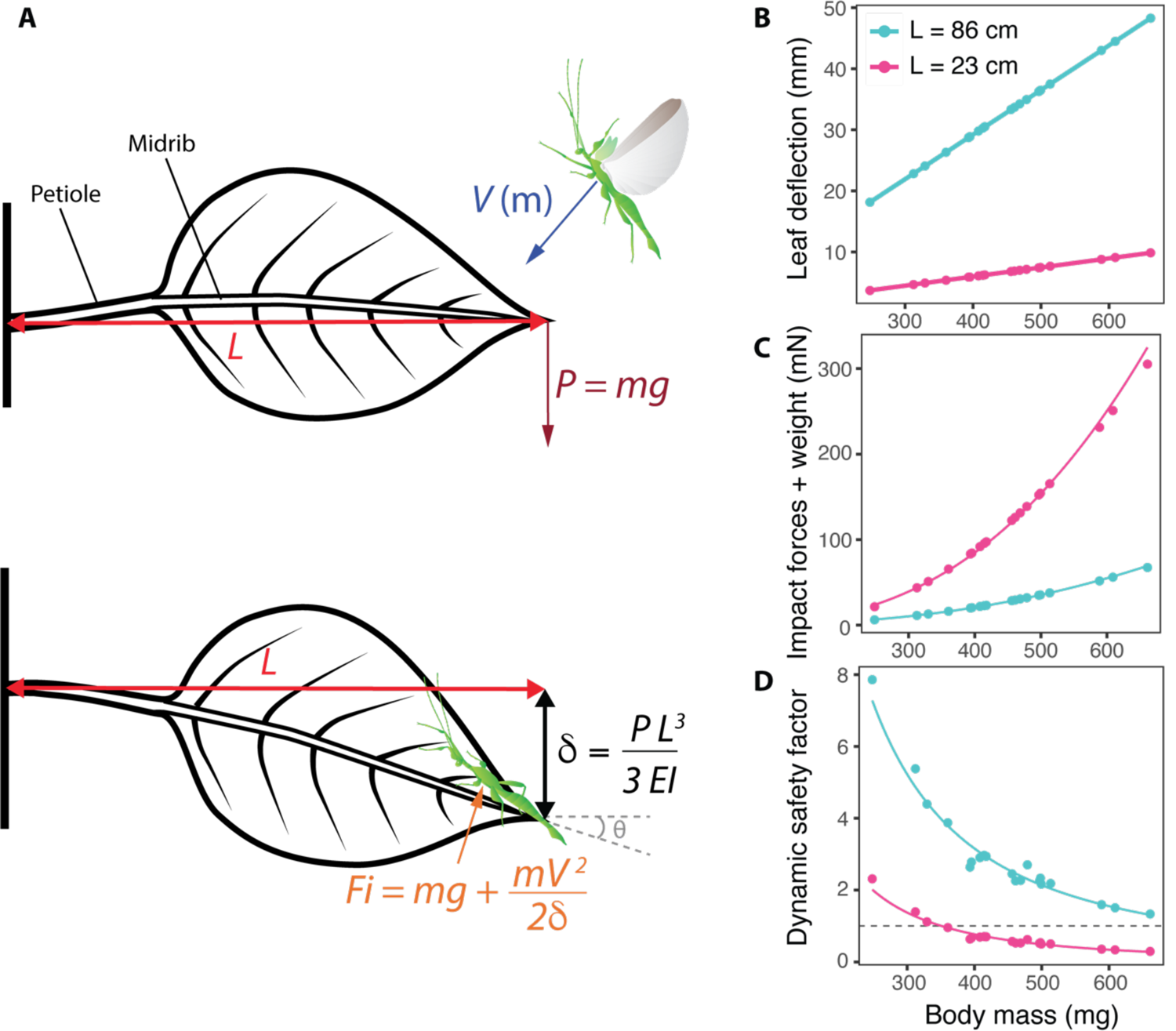
(A) Diagram of a male leaf insect landing on a leaf, including calculations used to estimate impact forces (F_i_) and dynamic safety factors (SF_dynamic_)– i.e., the ratio of frictional attachment forces and F_i_. The male lands with a landing velocity V which depends on the body mass of the insect (m). The terminal load on the leaf (P) will be equal to the weight of the insect – i.e., body mass (m) multiplied by the acceleration due to gravity (g). The subsequent deflection of the leaf (*δ*) depends on P, the length of the leaf (L) and the leaf flexural stiffness (EI). The impact force experienced by the male leaf insect (F_i_) will be determined by P, the kinetic energy of the insect (mV^2^/2) and *δ*. See the text for more detailed information. Relationships between male body mass (m) and *δ* (**B**), F_i_ (**C**) and SF_dynamic_ (**D**). Results for large leaves (L= 86cm) are represented in blue, small leaves (L=23cm) in pink. The dashed line in (**D**) represent a safety factor of one.

**Table 3:** Tests of the effect of body mass and leaf size on landing impact forces and dynamic safety factors in males. Type-I ANCOVAs were performed. Scaling exponents β and the corresponding 95% confidence intervals are shown for small and large leaves separately. Significant effects (i.e., p <0.05) are bolded.

| Response variable<br>(log<sub>10</sub> transformed) | Explanatory<br>variables | F | df1 | df2 | P | Slope $\beta$<br>[95% CI] |
| --- | --- | --- | --- | --- | --- | --- |
| Impact forces + weight | log<sub>10</sub>(body mass) | **20541** | **1** | **36** | **<0.001** |  |
|  | leaf size | **31459** | **1** | **36** | **<0.001** | Small leaf: 2.67 [2.6, 2.74] |
|  | interaction | **55.7** | **1** | **36** | **<0.001** | Large leaf: 2.41 [2.38, 2.43] |
| Dynamic friction safety factor | log<sub>10</sub>(body mass) | **892** | **1** | **36** | **<0.001** |  |
|  | leaf size | **2485** | **1** | **36** | **<0.001** | Small leaf: -2.02 [-2.21, -1.82] |
|  | interaction | **4.40** | **1** | **36** | **0.043** | Large leaf: -1.75 [-1.92, -1.57] |

### Body size increases the cost of flight

To further understand how body size and shape affect the costs of flight and consequently flight performance, we generated 3D models of the bodies of males of varying size and abdominal shape and estimated body lift and drag during steady horizontal flight using computational fluid dynamics (CFD, see methods) (Crandell et al., 2019; Goyens et al., 2015; Troelsen et al., 2019). CFD simulations showed that the male’s flattened abdomen produces a wide region of low-pressure behind the insect, whose size largely depends on its shape (Fig. 5). The males’ poorly streamlined bodies create high drag coefficients (1.38 < C_D_ < 1.55, Fig. 6C) and only nominal lift coefficients (0.85 < C_L_ < 1.03, Fig. 5D) resulting in relatively low lift to drag ratios (0.56 < L/D < 0.74, Fig. 6B). Drag must be matched by thrust provided by the wings, while body lift adds to weight support provided by the wings to balance body weight (Fig. 6A).

**Figure 5:**
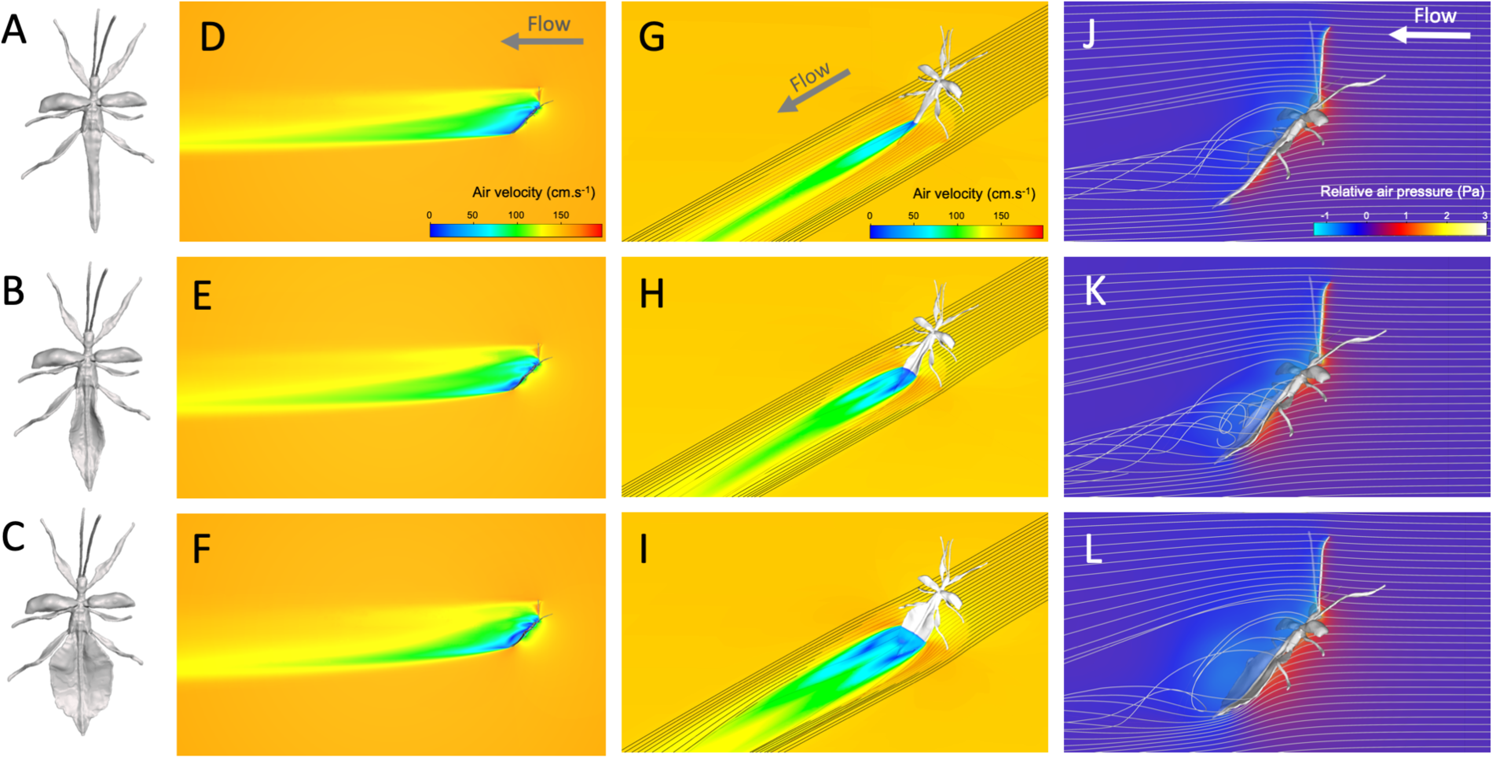
3D models and computational fluid dynamics simulations. Dorsal view of phasmid models with no (**A**), natural (**B**) and widened (**C**) ‘leaf-like’ abdominal expansions. Air velocity in the mid-sagittal plane (**D-F**), air velocity and streamlines in a horizontal transverse plane (**G-I**), pressure (relative to ambient air static pressure) and streamlines in the mid-sagittal plane (**J-L**) around the bodies of males with no (**D, G, J**), natural (**E, H, K**) and widened (**F, I, L**) abdominal expansions.

**Figure 6:**
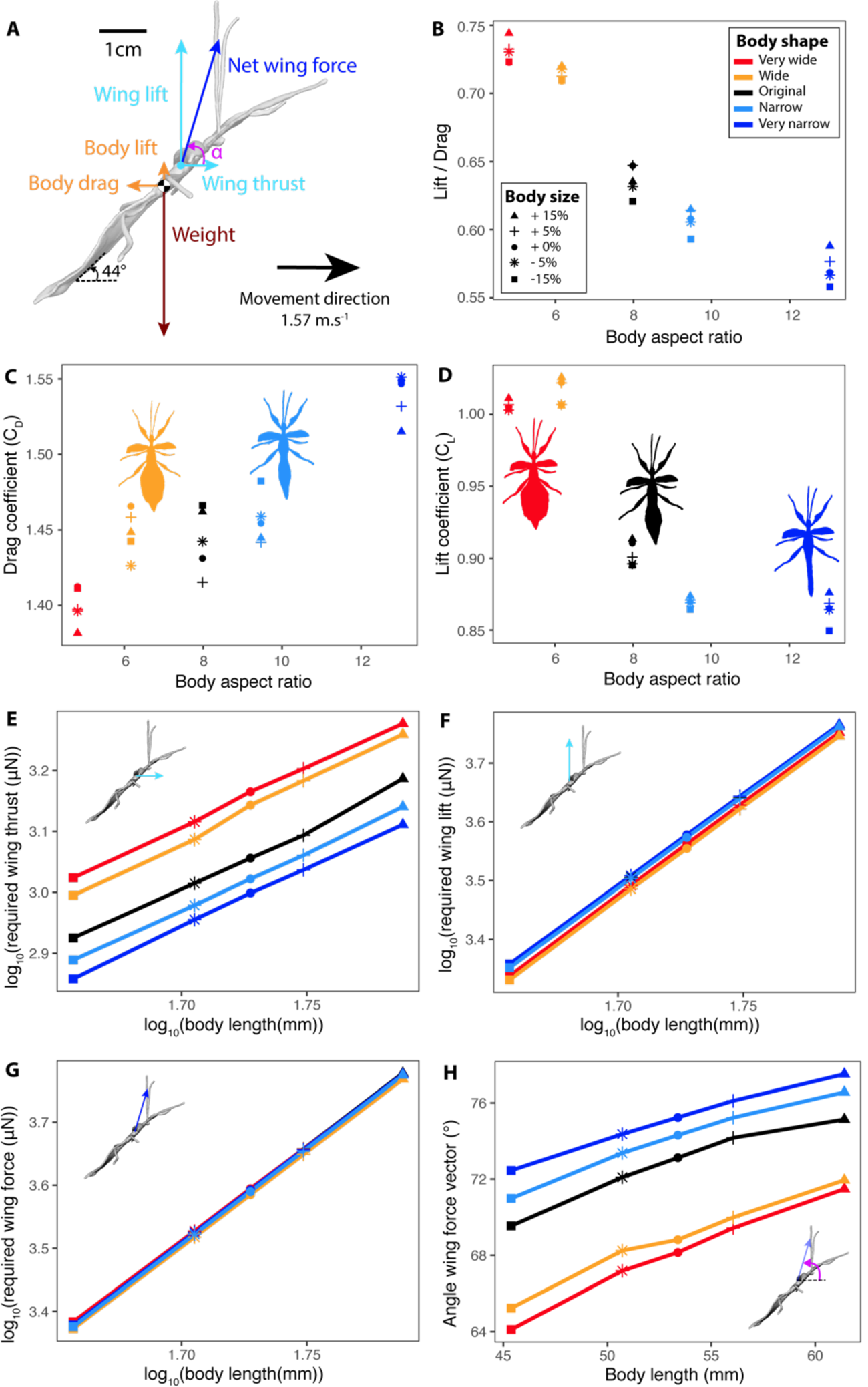
(A) Diagram of the forces applying upon a flying male leaf insect. Orange: aerodynamic forces estimated by the CFD simulations. Light blue: vertical and horizontal wing forces that must be generated to maintain a steady and horizontal trajectory. Dark blue: sum of the two wing force vectors. Pink: angle of the net wing force relative to horizontal. Lift to drag ratio (**B**), drag (**C**) and lift (**D**) coefficients as a function of model aspect ratio as estimated from the CFD simulations. Log-transformed wing thrust (**E**), lift (**F**) and net force (**G**) needed to maintain a steady flight as a function of log-transformed model body length. (**H**) Angle of the net required wing force relative to horizontal as a function of model body length. Colors in **B-H** correspond to different abdominal shapes of the models and point shapes correspond to different body sizes relative to the original model.

We found that larger males must provide greater wing thrust and weight support forces and therefore a higher resultant wing force relative to smaller males to maintain a steady horizontal flight (Fig.6E-G, Table 4). The angle of the net wing force vector becomes more vertical with increasing body size (Fig. 6H, Table 4): larger males need to provide more weight support relative to thrust to stay in the air, suggesting that balancing weight becomes relatively more important than offsetting drag as body size increases.

**Table 4:** Results of type I ANOVA from linear models contrasting the effects of body length and body aspect ratio on various aerodynamic variables. Significant effects (i.e., p <0.05) are bolded.

| Response variable | Explanatory variables | F | df1 | df2 | P |
| --- | --- | --- | --- | --- | --- |
| Lift to drag ratio (L/D) | Body length | 2.72 | 1 | 22 | 0.11 |
|  | **Body aspect ratio** | **221.7** | **1** | **22** | **<0.001** |
| Drag coefficient ($C_D$) | Body length | 2.42 | 1 | 22 | 0.13 |
|  | **Body aspect ratio** | **103.1** | **1** | **22** | **<0.001** |
| Lift coefficient ($C_L$) | Body length | 0.60 | 1 | 22 | 0.45 |
|  | **Body aspect ratio** | **68.3** | **1** | **22** | **<0.001** |
| $\log_{10}$ required wing thrust | **$\log_{10}(\text{body length})$** | **727.4** | **1** | **22** | **<0.001** |
|  | **$\log_{10}(\text{body aspect ratio})$** | **388.0** | **1** | **22** | **<0.001** |
| $\log_{10}$ required wing lift | **$\log_{10}(\text{body length})$** | **16911** | **1** | **22** | **<0.001** |
|  | **$\log_{10}(\text{body aspect ratio})$** | **46.3** | **1** | **22** | **<0.001** |
| $\log_{10}$ net required wing force | **$\log_{10}(\text{body length})$** | **32275** | **1** | **22** | **<0.001** |
| | $\log_{10}(\text{body aspect ratio})$ | 0.08 | 1 | 22 | 0.78 |
| Net wing force vector angle | **Body length** | **71.9** | **1** | **22** | **<0.001** |
|  | **Body aspect ratio** | **122.9** | **1** | **22** | **<0.001** |

### The power required for flight increases with size faster than the available muscle power

Flight performance depends on the power available (the maximum amount of mechanical energy that can be provided by the flight muscles per unit of time, P_a_) and on the power required for flight (the amount of mechanical energy required to fly per unit of time, P_r_) (Ellington, 1991). We estimated the scaling exponents of these two variables with BL using our empirical and modelling data to uncover how the difference between them (Δ*P*), which represents the excess power available for demanding aerial activities, varies with size. We show that P_a_ increases with body size with a scaling exponent β_3(_= 2, as expected under isometry. In contrast, P_r_ increases with body size with a scaling exponent β_3”_= 5.5 where isometry predicts β = 3.5 (see methods). Consequently, Δ*P* (P_a_ – P_r_) decreases with increasing body size more rapidly than would be expected under isometry, likely accounting for the reduced flight performance seen in larger males. Combined, our results suggest that selection for both flight and landing/attachment performance may help explain the relatively small size of males in this species.

### Body shape affects lift:drag ratio but there is a tradeoff with weight

Our CFD simulations further revealed that flying males with wider abdomens generate more lift relative to drag, have a lower C_D_, and a higher C_L_ (Fig. 6B-D, Table 4). As expected, wider males must provide higher wing thrust to offset the added drag in flight (Fig. 6E, Table 3). However, for a given body length, relatively wider abdomens generate more lift, even after accounting for the added weight (Fig. 6F, Table 4). Consequently, wider males must provide the same net amount of wing force to maintain themselves in the air as slenderer males (Fig. 6G, Table 4). The added weight and drag caused by the abdominal extensions were balanced by the additional lift they provided to the insect (Fig. 6H, Table 4). Thus, contrary to our predictions, our results suggest that abdominal shape does not affect the cost of flight and that selection for flight performance does not explain the relatively slenderer body shape of males in this species.

## Discussion

As leaf masqueraders, leaf insects (Phylliidae) display some of the most extreme abdominal morphologies of the insect world, and sexual size and shape dimorphisms so spectacular that taxonomists have had difficulty associating males and females of the same species (Cumming et al., 2020c). Using these organisms, we provide rigorous support for the often-assumed hypothesis that selection for locomotor performance favors small body sizes in males, contributing to the evolution of extreme sexual dimorphism in insects with scramble competition mating systems.

We show that small males have greater agility in the air, as they were able to stabilize their body angle faster after falling (Dudley, 2002), and they were better able to climb in flight than larger males. Flight is most likely essential to the mate searching performance of males, as females are typically scattered in the rainforest canopy (Herberstein et al., 2017). A decreased agility in the air and a lower ability to ascend during flight may be detrimental in that regard. Indeed, lost height may only be regained by walking up branches or trunks and, compared with flying, walking is much slower and has a higher cost of transport --i.e., a higher energy expenditure to move a given distance (Dudley et al., 2007; Schmidt-Nielsen, 1972). Caution is warranted when interpreting our results, though, as pitch control is only one aspect of maneuvering; roll and yaw likely also contribute (Cheng et al., 2016). Our computational fluid dynamic models further revealed that providing weight support becomes relatively more important than generating thrust in larger individuals, thereby highlighting the critical role of body weight for the flight performance of these insects. We estimated that the mechanical power required for these males to fly steadily and horizontally (P_r_) increased at a faster rate than the power available from the flight muscles (P_a_), and even faster than predicted under isometry. The reduction of flight performance (agility and climb ability) with body size seen in leaf insects is therefore likely to result from the decrease of ΔP (=P_a_–P_r_), which represents the excess power available for more demanding aerial activities such as maneuvers and climbing in air (Ellington, 1991; Norberg and Norberg, 2012).

Not only were large males relatively poor flyers, they also were more likely to detach and fall when landing. The largest males had the lowest dynamic safety factors for frictional attachment forces, which are fundamental for accommodating impact forces when landing (Higham et al., 2017). Indeed, we estimated that the largest males were likely to slip when landing on small and stiff leaves (i.e., SF_dynamic_ < 1). High attachment safety factors are crucial for canopy insects to avoid falling, which can lead to injury (Büscher and Gorb, 2017; Schmitt et al., 2018) and predator exposure (Dudley et al., 2007). For searching males, it can also mean losing track of a female. We found that male leaf insects have specialized, ridged surfaces on the pads of their tarsi which are not present in females (Figure 2B-C), and which are probably adapted to gripping a broad range of plant surfaces (Büscher et al., 2019; Büscher and Gorb, 2019; Bußhardt et al., 2012; Gorb and Scherge, 2000; Varenberg and Gorb, 2007, 2009). Flying males are likely to be confronted with unpredictable surfaces (e.g., branches, leaves) when walking and landing, and may benefit from generalist tarsal pads that adhere securely to a range of textured surfaces.

In contrast, females move very little in the canopy and are strongly associated with the leaves of their food plants. Females likely use their claws and attachment pads on a narrower range of surfaces and for a narrower range of tasks -- primarily as anchors as they hang upside down from smooth leaf cuticle. In this context, adhesion (as opposed to friction) forces are essential. In stick insects, arolia are shear-sensitive attachment pads providing most of the adhesion (Büscher and Gorb, 2019; Labonte and Federle, 2013). Consistently, female leaf insects have larger arolia relative to body size than males and consequently produce larger adhesion forces. Nevertheless, as females weigh much more than males, their static safety factors for both adhesion and friction forces were still relatively lower than those of males.

Male and female leaf insects also exhibit a spectacular interspecific variation in body shape related to leaf mimicry (48–52). This variation is likely driven by the coevolution between the insect appearance and its host plants. As exemplified by *P. philippinicum*, flight-capable males generally have more elongated body shapes than their respective flightless females and relatively reduced abdominal leaf-like lobes. Our CFD models showed that despite the poorly streamlined body shapes of males and their high drag coefficients (C_D_>1), wider abdomens do not incur significantly higher aerodynamic costs, contrary to our prediction. Wider abdomens are comparatively heavier and generate more drag, but they also generate relatively more lift resulting in a similar net cost of flight for thin or wide males. Therefore, flight performance does not seem to constrain male body shape the way that it does body size, and may have provided male leaf insects the freedom to evolve a variety of body shapes (Brock et al., 2020; Conle et al., 2008; Cumming et al., 2020a, 2020b, 2018).

## Materials and Methods

### Study animals

A first breeding population of *P. philippinicum* was obtained from the Audubon Insectarium in New Orleans, Louisiana, USA and shipped to the University of Montana, Missoula, Montana, USA. The insects were housed in a transparent plastic container (50×40×60cm) at 22°C, on 12h:12h light:dark cycles, sprayed with water daily (RH = 50 – 80%), and fed fresh *Rubus idaeus* leaves *ad libitum*. This population was used to investigate morphological scaling relationships and conduct the flight experiments described below.

A second culture stock of *P. philippinicum* was obtained from Kirsten Weibert (Jena, Germany) and captive bred in the department of Functional Morphology and Biomechanics at Kiel University, Germany. The specimens were kept in a large glass cage with proper ventilation at 20-22°C (RH = 50 – 80%; 16h:8h light:dark cycle) and fed with fresh blackberry (*Rubus* sp.) and common oak (*Quercus robur* L.) leaves *ad libitum*. This population was used to investigate attachment pad morphology and attachment forces.

### Sexual dimorphism and scaling relationships

Photographs of the animals in dorsal view were taken using a DSLR camera (EOS 600D, Canon Inc., Tokyo, Japan). Using ImageJ software (v.1.52k) (Schneider et al., 2012), we measured body length (BL, mm), body area (mm^2^), body circularity 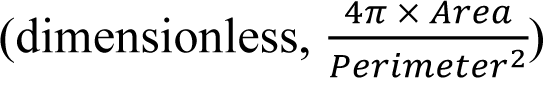 body aspect ratio 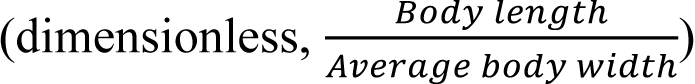 mean antenna length (mm), mean front femur length (mm), and total wing area (mm^2^, including both forewings for females or both hindwings for males) in 25 adult males and 19 adult females (Fig. S1). Antenna length could not be measured in seven males and three females because they were missing flagellomeres on both antennae. Wet body mass (g, measured immediately after flight trials) was obtained using an analytical balance (ME54TE/00, Mettler Toledo, Columbus, OH, USA). We calculated wing loading (N.m^-2^) as wet body mass multiplied by gravitational acceleration (g=9.81 m.s^-2^) and divided by total wing area. Male flight muscle mass (mg) was obtained by dissecting the muscles out of the metathorax of freshly dead males (n = 23) drying them at 70°C for 24h and weighing them with a more accurate analytical balance (UMT2, Mettler Toledo, Columbus, OH, USA).

### Male flight performance

To evaluate male flight performance, here defined as a righting maneuver that required sustained control over long-axis body angle and climb ability, we dropped adult males in the air and recorded their flight trajectories in 2D (Video S1-2). Working in Missoula, MT (elevation 978 meters above sea level, average air density = 1.07 kg.m^-3^), adult males (N=16, 0.43 ± 0.006 g) from our American culture population were held by the thorax and dropped by the experimenter at a horizontal body angle from a constant height above a floor (1.5m). A 2m-tall slab of ponderosa pine (*Pinus ponderosa*) with natural bark was placed vertically in front of the animal, 2m away, to serve as a target and landing site (Fig. S3). The experimental room was largely featureless with white walls and at 26°C. The floor was covered with thick blankets to avoid injuries if crashing. We recorded the flight of the insects using a high-speed video camera (Photron FASTCAM SA-3, Photron USA, San Diego, CA, USA) sampling at 500 fps with a shutter speed of 1/5000s and 1024×1024 pixel resolution (Photron PFV v.3.20). Because the insects were induced to fly in a trajectory parallel to the plane of the imaging sensor (deviations < 10°), we analyzed the trajectories in two dimensions (Fig. S3, S4A). Pixels were scaled to metric coordinates using a 50cm bar held horizontally at the same level of the flight trajectories. We recorded a clay ball in free fall to calibrate the vertical direction. Average vertical ball acceleration was 9.816 ± 0.14 m.s^-2^, less than 1% different from gravitational acceleration (9.805 m.s^-2^) and the camera was oriented to gravity so that the vertical ball drop direction of acceleration was always <5% of 90°. Each insect was dropped several times (maximum five times) until we obtained two straight trajectories per animal. A resting period of approximately 20min was left in between each flight to allow recovery. Males were frozen-killed at −80°C just after the experiments. After estimating the relative position of their center of mass (see below), males were pinned with their wings fully extended for morphometrics.

Video digitization was done by tracking morphological landmarks using the open source video analysis tool *DLTdv5* by T. Hedrick implemented in MATLAB (R2016b, MathWorks, Natick, MA, USA) (Hedrick, 2008) and the obtained data was analyzed using R (v 3.6.1)(R Core Team, 2019). In *DLTdv5*, We used autotracking mode (predictor tool: extended Kalman) and manual tracking when the autotracking mode was unreliable. We marked the position of the head and of the terminal abdominal segment on each frame. Body pitch (°) was calculated for every frame by calculating the angle between the horizontal and the line linking the position of the head and that of the terminal segment (Fig. S4B). Typically, in the first phase of the fall (free fall), body pitch decreases (i.e., the insect rotates forward, eventually diving head first), before the insect opens its wings (t_1_) and actively corrects (t_2_) and stabilizes its body pitch (phase 3) (Fig. S4B). Body pitch was smoothed using a Savitzky-Golay filter with a polynomial order of 3 and a window size of 71 (*sgolayfilt*: ‘signal’, *function*:’R package’). The beginning of phase 2 (t_1_) was determined as the time corresponding to the minimal body pitch (Fig. S4B). The end of phase 2 (t_2_) corresponded to the time when the insect’s pitch stabilized – *i.e.*, when the rotational velocity (°.s^-1^) of body pitch reached a local minimum after a large peak corresponding to phase 2 (Fig. S4B). We calculated the average rotational velocity during phase 2 (ω) as:

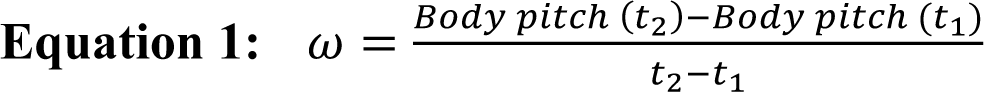

We used ω as to quantify torsional agility -- an important aspect of maneuverability -- as it reflects how fast the animal can rotate to correct its body pitch in the air from a free falling, head first, position to a stable flight body pitch. This correction occurred over several wingbeats and was therefore distinct from within and among-wingbeat oscillations which could have indicated a lack of longitudinal stability (Taylor and Thomas, 2003). The 2D position of the body center of mass was estimated using images taken in lateral view of the males (freshly dead) orthogonally balancing on a horizontal razorblade. Its position relative to the two landmarks (i.e., head and terminal segment) was then calculated which enabled us to define the center of mass of the individual on each flight video. Trajectories of the center of mass were analyzed using the package “trajr” in R (McLean and Skowron Volponi, 2018). Raw trajectories were smoothed using a Savitzky-Golay filter with a polynomial order of 3 and a window size of 31 (*TrajSmoothSG*:‘trajr’; Fig. S4A). Horizontal, vertical and composite velocities and accelerations were then computed on the smoothed trajectories (*TrajDerivatives*:‘trajr’; Fig. S4C-E). For each trial, we defined transient and steady states for both vertical and horizontal velocities (Fig. S4D-E). In both cases, the transient state corresponded to the free fall and maneuver of the insect in the air during which body velocity greatly varied. The steady state started when velocity stabilized and acceleration started oscillating around 0 m.s^-2^ (Fig. S4D-E). We extracted the mean vertical and horizontal velocity (m.s^-1^) during their respective steady states. These measures were used to quantify the capacity of the insect to fly forward and ascend. On the videos, the position of the tip of the wing closest to the camera was also manually marked on frames corresponding to the end of upstroke and downstroke as this was sufficient to measure wing beat frequency (Hz) and stroke amplitude (°). Average wing beat frequency was calculated after the animal reached a stable body pitch (*i.e.,* after t_2_; Fig. S4B). For each animal, we measured wing length and the position of the attachment of the wings relative to the head and tip of the abdomen using photographs and ImageJ. We then determined the position of the wing attachment point on each frame on the videos using this relative position between our two landmarks. The amplitude of wing strokes was calculated using trigonometry (Fig. S5).

### Adhesive pads and substrate attachment performance

We used scanning electron microscopy (SEM) to observe the tarsi of the metathoracic leg of adult males and females, measure attachment pad areas, and describe the microstructures on these pads. Tarsi of the right metathoracic leg were cut off from 20 adult males and 20 adult females and fixed in 2.5 % glutaraldehyde in PBS buffer for 24 h on ice on a shaker, dried in an ascending alcohol series, critical-point dried and sputter-coated with a 10nm layer of gold-palladium. To obtain overview images, we used a rotatable specimen holder (Pohl, 2010) and the scanning electron microscope (SEM) Hitachi TM3000 (Hitachi High-technologies Corp., Tokyo, Japan). The micrographs for visualization and measurements were taken at an acceleration voltage of 15kV. The attachment microstructures on the tarsi of both sexes were further examined using the SEM Hitachi S4800 (Hitachi High-Technologies Corp., Tokyo, Japan) at 7 kV of acceleration voltage. Processing of the raw micrographs and measurements of projected attachment pad area (mm^2^) -- i.e., the surface area of the tarsus specialized for adhesion and friction (Bullock et al., 2008; Labonte et al., 2016) -- were done using Photoshop CS6 (Adobe Systems Inc., San José, CA, USA).

To measure attachment forces (mN) in both pull-off (adhesion) and traction (friction) directions, we used 20 adult males (Mean ± S.D.= 0.46 ± 0.02g) and 20 adult females (5.22 ± 0.31g). A horsehair was glued to the metanotum of each insect and then attached to a 100g force transducer (FORT100, World Precision Instruments, Sarasota, USA, linearity error: <0.1%, resolution: 0.01%), connected to a BIOPAC model MP100 and TCI-102 system (BIOPAC Systems, Inc., Goleta, CA, USA), and mounted on a motorized micromanipulator (DC 3001R, World Precision Instruments Inc.). Maximum adhesion forces were recorded by vertically pulling the insects off a horizontal glass plate until they detached from the glass plate (Büscher and Gorb, 2019; Wohlfart et al., 2014). The micromanipulator was moved upwards with a speed of 200 µm/s at a step size of 10 µm until the specimen was detached from the surface as indicated by an instantaneous drop in force. Maximum friction forces were recorded by horizontally pulling the insects backwards with the same retraction velocities as above, until detaching them from the glass plate (Büscher and Gorb, 2019; Wolff and Gorb, 2012). A glass plate was used as the substrate for the attachment force measurements, to eliminate mechanical interlocking of the claws with surface irregularities of rough substrates, or penetration of soft substrates. As glass is smooth on the microscopical level, this substrate enables estimation of the traction and pull-off performance of the attachment pads themselves on a standardized level, without the influence of substrate irregularities. Force-time curves were obtained using Acqknowledge 3.7.0 (BIOPAC Systems Inc., Goleta, CA, USA) and the maximum peaks were extracted as maximum adhesion forces, or maximum friction forces respectively. Each of the 20 males and females were measured three times in both directions on a glass plate and the average of the three measurements was used as the individual maximum adhesion/friction force. The order of the individuals was randomized and the substrate was cleaned between every measurement. The experiments were conducted at 20-23°C and 50-60% relative humidity.

Static safety factors 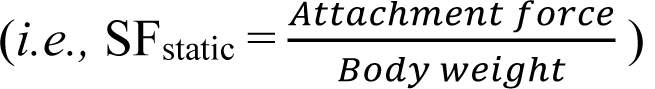 were computed for each individual. Following the methods of Higham et al., 2017, we estimated impact forces (F_i_, in N) during landing for each male to compute dynamic safety factors 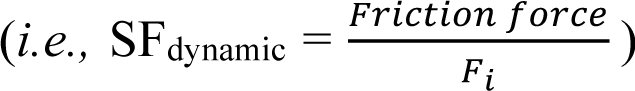 We assumed that males would stop immediately after landing on the tip of a leaf without slippage. F_i_ was calculated using the work-energy principle:

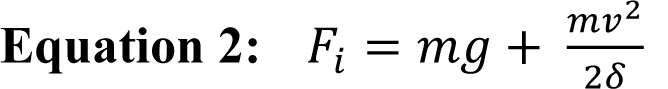

where m = mass of the insect (kg), g = acceleration of gravity (9.81 m.s^-2^), v = landing speed of the insect (m.s^-1^) and δ = deflection of the leaf (m) (Fig. 4).

We used a simplified model of leaf to estimate a range of values for the deflection of a leaf upon landing of a male leaf insect and explore potential values of SF_dynamic_. The leaf and petiole were considered as a uniform cantilever beam with impact forces applying at the tip at length L. We considered the leaf to be initially horizontal. Deflection was calculated as a function of the weight of the insect. Given the relatively light weight of male leaf insects, and in contrast with Higham et al., 2017, who were considering geckos (i.e., roughly 28 times heavier), we only accounted for small leaf deflections (deflection angle θ < 5°) and therefore used the following equation to estimate the deflection of the tip of the leaf δ (Goodno and Gere, 2018):

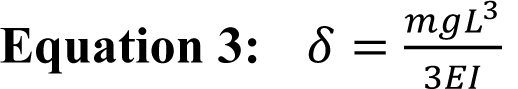

where L is the total length of the leaf (m) and EI is its flexural stiffness (N.m^2^). We used the range of values estimated by Higham et al., 2017, for leaf length and corresponding EI (i.e., L_1_ = 23 cm, EI_1_ = 2.67 × 10^-3^ N.m^2^ and L_2_ = 86 cm, EI_2_ = 28.48 × 10^-3^ N.m^2^) to explore the possible range of SF_dynamic_.

Finally, landing speed was estimated as a function of the body mass of the individual. Using our experimental flight videos (see above), we extracted the instantaneous speed of the experimental males right before making contact with the wood slab (i.e., the landing target) and built a linear regression between landing speed and body mass, including individual ID as a random factor (*lmer*: ‘lme4’, Bates et al., 2015). We found a significant positive effect of body mass on landing speed (see results) and used the fixed-effect estimates (intercept and slope) from this model to predict the landing speed of the males for which we measured attachment forces, given their body mass. This estimated landing speed was used in equation 2.

### Computational fluid dynamic simulations

To investigate how lift and drag produced by the insect were affected by size and shape, we generated 3D models of males of varying size and shape and estimated these forces during a steady and horizontal flight using computational fluid dynamics (CFD). We created a reference 3D surface model of an adult male body using photogrammetry. We pinned the body of a dead specimen in a flight posture (i.e., legs extended, forewings opened perpendicularly, antennae oriented 50° up, hindwings removed). We did not model flapping aerodynamics as fully integrating the complex three dimensional trajectories and aeroelastic deformations of the wings into a CFD model (e.g., Young et al., 2009) was beyond the scope of this study. The aerodynamic interactions of the flapping wings with the body of insects in slow flight has been shown to be negligible (∼5%) in (Liang and Sun, 2013). Once dried, the individual was vertically mounted onto a pin on a custom-made turntable. 2D images using a DSLR camera (EOS 600D, Canon Inc., Tokyo, Japan) equipped with a macro lens (Canon EF 100mm f/2.8 Macro USM), were then obtained from 100 different orientations (Fig. S6). The 3D model was then reconstructed from these multiple images using Autodesk ReCap Pro 2019 (v5.0.4.17, Autodesk Inc., San Rafael, CA, USA) and subsequently smoothed and rewrapped using Autodesk Meshmixer 2017 (v11.5.474, Solid accuracy: 402, cell size: 0.202, density: 219, offset: 0.25, min thickness: 0.14mm). From this reference model, we built four additional and artificial models using the “Move” tool in Meshmixer to either manually extend or shrink the abdominal lobes and therefore manipulate abdominal shape. These artificial shapes purposely spanned a wider range of body aspect ratios than the one found in actual males *P. philippinicum* (male natural range: 4.89 - 6.42, female natural range: 2.28 - 2.84, model range: 2.28 - 9.47). The model with the lowest aspect ratio displayed a female-like abdominal shape while the model with the highest aspect ratio had no abdominal expansions. The models were further scaled to a body length of 53 mm (mean male BL in our Montana population = 52.6 +/- 0.25mm) using Autodesk fusion 360 (v2.0.8335). From each of these five meshes, we created four additional models (25 models in total) respectively scaled to a factor 0.85, 0.95, 1.05 and 1.15. The insect models were tilted at a 44° body pitch in our control volume. This angle was determined using our flight experiments and a LMM with stable body pitch as the response variable, horizontal and vertical body velocity as main effects and individual ID as a random factor. Using the parameters estimated by this model, we predicted a body pitch of 43.9° for a vertical velocity of zero and a mean horizontal velocity of 1.57 m.s^-1^ (i.e., the average horizontal velocity calculated from our flight trials, after the animal had stabilized its body pitch: 157 +/- 6.8 cm.s^-1^).

We constructed a control volume around these body meshes in Autodesk CFD 2019 (v19.2), that provided numerical solutions to the Reynolds-averaged Navier-Stokes equations (Rahman, 2017; Troelsen et al., 2019). A fluid volume was built around the mesh with walls far enough from the model mesh to avoid any reflection effects (1×0.5×0.5m) (Fig. S7A). The fluid was assigned the default properties of air in CFD 2019 (density at sea level = 1.205 kg.m^-3^, viscosity at 20°C = 18.2 μPa.s^-1^). The phasmid models were then assigned properties of hardwood which, for mass-less and stationary models, should have no impact on results. The input flow on the anterior end of the control volume was set to 1.57 m.s^-1^. We applied a zero-pressure condition on the opposing end of the volume. A slip/symmetry condition was applied to all other fluid boundaries. We automatically meshed the domain around the phasmid model, applied a surface refinement, and locally defined a non-uniform mesh refinement region (0.7×0.2×0.2m) with a mesh size reduced to 75%, around and behind the model to better capture the resulting wake. We ran steady-state simulations using the turbulence model k-epsilon. The maximum number of iterations was set to 3,000 although the simulations were stopped when they reached convergence according to the default convergence detection parameters of the CFD software (mean = 849 ± 50 interactions). The adaptive meshing tool was used to insure mesh optimization for our models and mesh independence of the results. The simulation was first run with the meshing parameters described previously. Then, the solution results of this simulation were automatically used to refine the mesh in high velocity gradient regions and rerun the simulation. We enabled the ‘flow angularity’ option to improve mesh resolution in areas with a lot of flow separation, the ‘free shear layers’ and ‘external flow’ options to refine the mesh in areas of strong velocity gradients. We ran 3 such cycles for each model. Final mesh sizes averaged 865,446 ± 74,771 nodes and 4,411,596 ± 379,780 elements (Fig. S7). Finally, to help evaluate the validity of our simulations, we placed a sphere with the same Reynold’s number as the one calculated for the original phasmid model (Re = 5558) in a similar control volume with the exact same settings as our insect simulations. We found a drag coefficient (Cd) of 0.643. This is very close to the value predicted from experimental data (Cd = 0.652) and which was determined using the equation 8.83 in Morrison, 2013.

The weight of the models (mN) with a non-modified abdominal shape (reference models) was determined from their BL using the linear regression built between male body mass and BL in our American population. To estimate the weight of the models with artificial abdominal shapes, we first measured their abdominal area relative to that of the reference model of identical BL and measured the average weight of the leaf-like abdominal expansions by cutting these extensions from 5 freshly frozen-killed males and measuring their areas and mass (89.7 ± 0.6 g.m^-2^). The CFD simulations estimated the aerodynamic forces (drag and lift, mN) that apply to the rigid insect body flying horizontally and steadily. We then calculated the forces the wings would need to provide to balance body drag and weight and maintain a steady and horizontal flight. Wing lift and thrust values, and the norm and angle of the sum of these two vectors (i.e., net wing force, NWF) were determined (Fig. 5A, equations S1 & S2). The net required wing forces derive from the mechanical work done by the flight muscle. Therefore, as in Goyens et al., 2015, we considered these forces as a proxy for the net mechanical cost of flight (i.e., the mechanical energy expenditure necessary per unit of distance covered). It should be noted that weight and aerodynamic forces acting on the body act on its center of mass while wing forces apply where the wings attach to the body (Fig. 5A). This discrepancy could create a potential torque that would need to be compensated to maintain a stable body angle, but, for simplicity, it is not further considered.

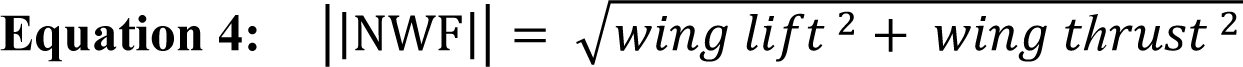

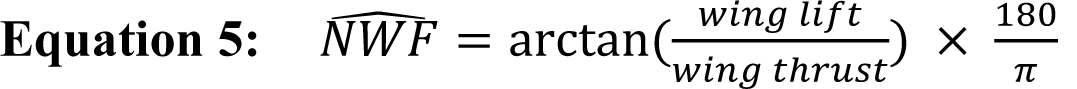

For each model, we measured the projected frontal area on a plane perpendicular to the air flow. We then calculated their coefficient of drag 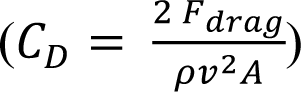 and lift 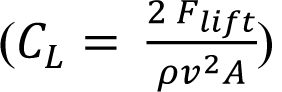 using the model frontal area (A), the mass density of air (= 1.20473 kg.m^-3^), the velocity of the insect (v =1.57 m.s^-1^) and the drag or lift force estimated from the CFD simulations. Lift to drag ratios 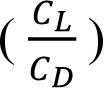 were also calculated for each model.

### Scaling of muscle power available and power required for flight

We estimated the scaling relationships of the power available (P_a_) and the power required for flight (P_r_) with body size using our empirical and modelling data. As leaf insects are slow flyers (average flight speed=1.70 ± 0.07 m.s^-1^, mean ± SE), P_r_ mostly corresponds to the induced power P_ind_ – i.e., the cost for producing lift (Biewener and Patek, 2018; Ellington, 1991). P_ind_ is the product of the net required force from the wings to maintain the animal in the air and of the induced velocity in the wake. Following Epting and Casey, 1973, we assumed that, under isometry, the induced velocity in the wake and weight-specific power required for slow flight should be proportional to the square root of wing disc loading (DL, N.m^-2^) --i.e., body weight (W_B_) divided by wing disc area (A_WD_). Wing disc area --i.e., the area swept out by the wing during a wing beat cycle and through which air is accelerated downward to develop lift force-- is determined by wing length (L_w_) and stroke amplitude (θ, °) (Equation S3).

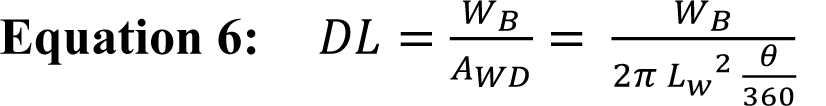

Thus, among geometrically similar animals, the power required for slow flight should scale as BL^7/2^ (Equation S4).

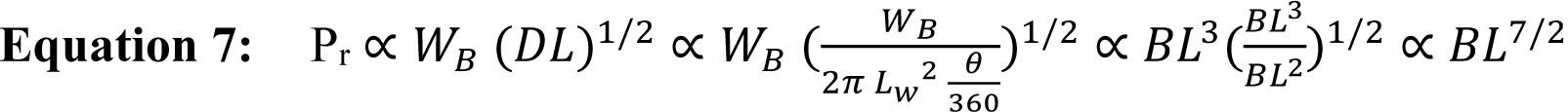

In slow flight, Pa is the product of the net wing force and of the tangential velocity of the tip of the wing --i.e., angular velocity (rad.s^-1^) × wing length (m)-- or the product of the muscle work --i.e., force distance of contraction-- and of flapping frequency (Hz). We accepted the assumption that, for geometrically and dynamically similar organisms, force is proportional to the cross-sectional area of the muscles, which scales as BL^2^, and distance of contraction scales as BL^1^. Thus, work scales as BL^3^ and is therefore directly proportional to W_B_ (Ellington, 1991; Hill, 1950). Flapping frequency is predicted to scale as BL^-1^ when the animal is using maximal or near-maximal effort (Greenewalt, 1975). Therefore, under isometry, P_a_ is expected to scale as BL^2^.

To empirically determine the scaling exponent of the tangential velocity of the tip of the wing (i.e., angular velocity (rad.s^-1^) × wing length (m)) and of wing disc loading (Eq.S3) with L, we built LMMs with log_10_ tangential velocity or log_10_ disc loading as the response variable, log_10_ BL as the fixed effect and male ID as a random factor. We similarly calculated the scaling exponent of the net required force from the wings with BL using our CFD models using a linear model. Tangential velocity of the tip of the wing did not significantly scale with BL in our flight experiments (β = −2.43 ± 1.60, *X*^2^= 2.46, df = 1, p = 0.12). The observed scaling exponent of flight muscle mass did not significantly differ from isometric expectations (Table S1). Coupling this with the aforementioned assumptions about muscle force, we estimated P_a_ scaled as BL^2^ in leaf insects, as expected under isometry. Net required force from the wings increased as BL^3^ in our CFD simulations (β = 3.01 ± 0.02, 95% CI=[2.98, 3.05]). Wing disc loading (DL) positively scaled with BL (*X*^2^ = 8.99, df=1, p=0.003). While, under isometry, we expected a scaling exponent of 1 (Equation 2), we found that DL increased disproportionately with BL (β = 4.96, 95% CI=[1.99,7.94]). This is a consequence of the reduced wing stroke amplitude seen in larger individuals (Table 2). As the induced velocity (V_i_) in the wake scales proportionately with the square root of wing disc loading, we estimated that P_r_ scaled more steeply with size than expected under isometry (β=5.48 VS 3.5) (Equation S5)

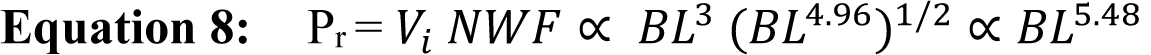

Consequently, ΔP (P_a_ – P_r_) decreased with body size more rapidly than would be expected under isometry.

### Statistical analyses

All statistical analyses were run in R version 3.6.1 (R Core Team, 2019). For all linear models, we systematically checked the normal distribution of the residuals and the absence of any specific patterns in their distribution.

We tested for sex differences in mean BL, body mass, body aspect ratio and antenna length using Wilcoxon-Mann-Whitney tests (*wilcox.test*: ‘stats’). To test for sex differences in the scaling relationships (i.e., in slope and intercept) of the various morphological traits, attachment and aerodynamic force measurements with BL, we built ordinary least square regressions (Kilmer and Rodríguez, 2017) including each log_10_-transformed trait as response variable and log_10_ BL, sex and their interaction as predictor variables (*lm*: ‘stats’). Type-I ANCOVAs were used to determine significance of the fixed effects (*anova*: ‘stats’). Departure from isometry which corresponds, on a log-log scale, to a slope of 1 for linear measurements, 2 for areas and 3 for masses (Schmidt-Nielsen, 1984), was tested using 95% confidence intervals (CI) around the estimated regression slopes (*confint*: ‘stats’). We similarly built linear models to investigate the effect of body mass, leaf size (i.e., small or large) and their interaction on landing impact forces in males and dynamic safety factors.

To test for effects of body size, wing size and body shape on male flight performance, we built linear mixed models (LMM) (*lmer*: ‘lme4’). Response variables were either rotational velocity (ω), mean vertical or horizontal velocity, wingbeat frequency, or wing stroke amplitude. Body mass, wing area and body aspect ratio were mean-centered and standardized (μ = 0 and σ = 1, *scale*: ‘base’) and were included as main fixed effects. Individual ID was included as a random factor to account for replications of each individual. Likelihood ratio tests were subsequently performed sequentially to assess the significance of the fixed effects (*anova*: ‘lme4’). For response variables significantly affected by body mass and wing area and to illustrate their combined effect, we built and plotted similar LMMs but with wing loading as the only fixed effect.

Following our CFD simulations, we tested for the effects of body size and shape on lift to drag ratio, C_D_, C_L_, required wing thrust, required wing lift, net required wing force and the angle of the net required wing force, using linear models including BL and body aspect ratio as explanatory variables. Sequential ANOVAs (type I) were subsequently performed to assess significance (*anova*: ‘stats’). In models including force estimates as response variables, variables were log_10_-transformed to compute scaling exponents as isometry predicts a power-law relationship between force and size (Schmidt-Nielsen, 1984).

## Acknowledgements

The authors thank Art Woods for use of lab computer and space and Camille Thomas-Bulle and Anthony Lapsansky for useful discussions.

## Data availability

Datasets and corresponding R scripts are publicly available on Figshare: Boisseau, Romain (2021): Boisseau et al._Phyllium locomotion 2021_data and scripts.zip. figshare. Dataset. https://doi.org/10.6084/m9.figshare.14852904.v1

---

**Table S1:**
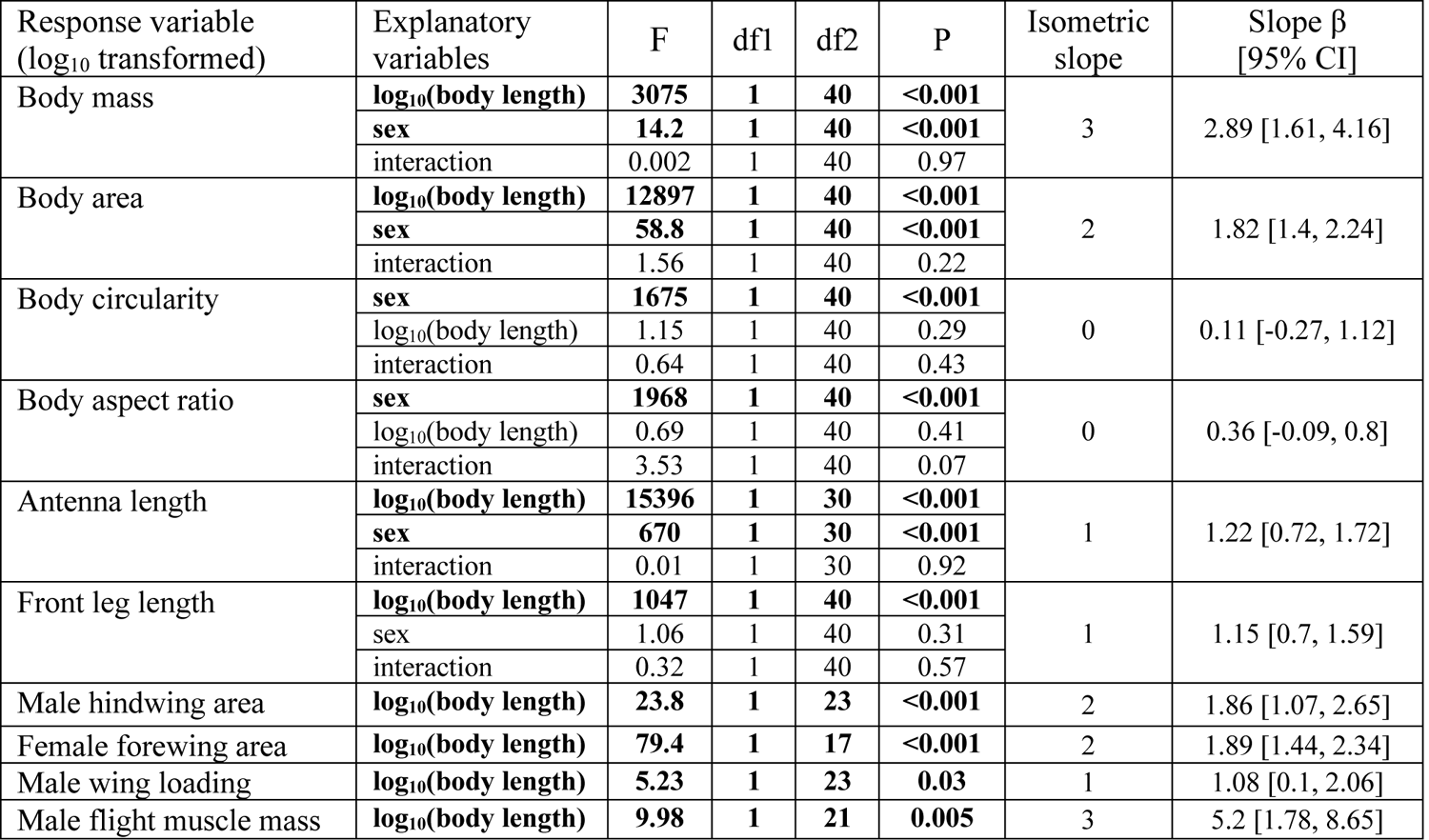
Results of type I ANOVA from linear models contrasting the differences between sexes in terms of scaling relationships between body size (body length) and body mass, area, circularity, aspect ratio, antenna length or front femur length. The scaling relationships for male hindwing area, female forewing area, male wing loading and male flight muscle mass with body size are also presented. Scaling exponents β and the corresponding 95% confidence intervals are shown in comparison to isometric expectations. Significant effects (i.e., p <0.05) are bolded.

| Response variable<br>(log<sub>10</sub> transformed) | Explanatory<br>variables | F | df1 | df2 | P | Isometric<br>slope | Slope $\beta$<br>[95% CI] |
| --- | --- | --- | --- | --- | --- | --- | --- |
| Body mass | **log<sub>10</sub>(body length)** | **3075** | **1** | **40** | **<0.001** | 3 | 2.89 [1.61, 4.16] |
|  | **sex** | **14.2** | **1** | **40** | **<0.001** |  |  |
|  | interaction | 0.002 | 1 | 40 | 0.97 |  |  |
| Body area | **log<sub>10</sub>(body length)** | **12897** | **1** | **40** | **<0.001** | 2 | 1.82 [1.4, 2.24] |
|  | **sex** | **58.8** | **1** | **40** | **<0.001** |  |  |
|  | interaction | 1.56 | 1 | 40 | 0.22 |  |  |
| Body circularity | **sex** | **1675** | **1** | **40** | **<0.001** | 0 | 0.11 [-0.27, 1.12] |
|  | log<sub>10</sub>(body length) | 1.15 | 1 | 40 | 0.29 |  |  |
|  | interaction | 0.64 | 1 | 40 | 0.43 |  |  |
| Body aspect ratio | **sex** | **1968** | **1** | **40** | **<0.001** | 0 | 0.36 [-0.09, 0.8] |
|  | log<sub>10</sub>(body length) | 0.69 | 1 | 40 | 0.41 |  |  |
|  | interaction | 3.53 | 1 | 40 | 0.07 |  |  |
| Antenna length | **log<sub>10</sub>(body length)** | **15396** | **1** | **30** | **<0.001** | 1 | 1.22 [0.72, 1.72] |
|  | **sex** | **670** | **1** | **30** | **<0.001** |  |  |
|  | interaction | 0.01 | 1 | 30 | 0.92 |  |  |
| Front leg length | **log<sub>10</sub>(body length)** | **1047** | **1** | **40** | **<0.001** | 1 | 1.15 [0.7, 1.59] |
|  | sex | 1.06 | 1 | 40 | 0.31 |  |  |
|  | interaction | 0.32 | 1 | 40 | 0.57 |  |  |
| Male hindwing area | **log<sub>10</sub>(body length)** | **23.8** | **1** | **23** | **<0.001** | 2 | 1.86 [1.07, 2.65] |
| Female forewing area | **log<sub>10</sub>(body length)** | **79.4** | **1** | **17** | **<0.001** | 2 | 1.89 [1.44, 2.34] |
| Male wing loading | **log<sub>10</sub>(body length)** | **5.23** | **1** | **23** | **0.03** | 1 | 1.08 [0.1, 2.06] |
| Male flight muscle mass | **log<sub>10</sub>(body length)** | **9.98** | **1** | **21** | **0.005** | 3 | 5.2 [1.78, 8.65] |

**Figure S1:**
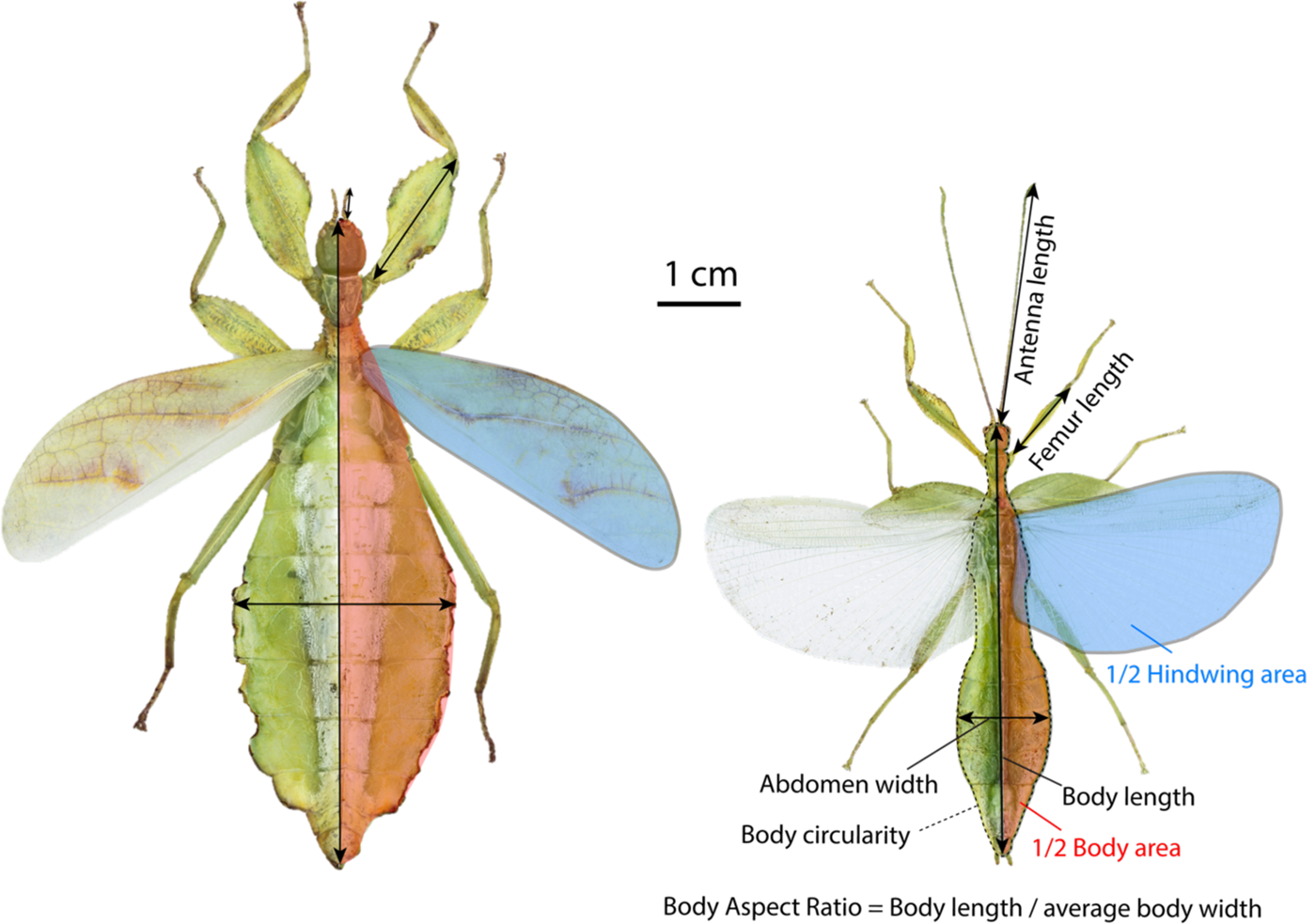
Morphological measurements for females (left) and males (right) *P. philippinicum*.

**Figure S2:**
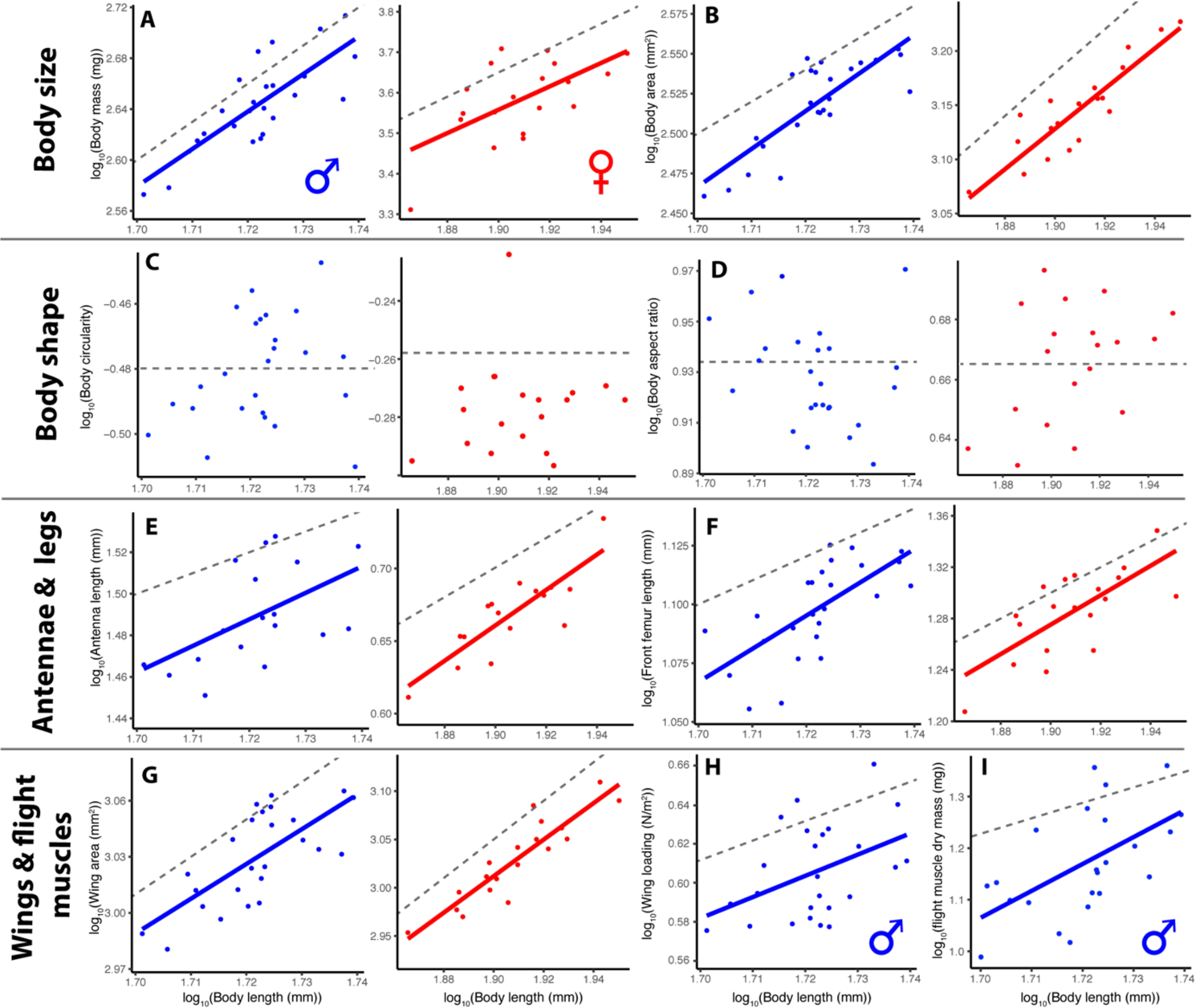
Scaling relationships between various morphological traits (body mass (**A**), body area (**B**), body circularity (**C**), body aspect ratio (**D**), antenna length (**E**), front femur length (**F**), total wing area (**G**), wing loading (**H**) and flight muscle dry mass (**I**)) and body length in males (blue) and females (red). Dashed lines show an isometric slope (arbitrary intercept). Wing area refers to hindwings for males, forewings for females. Wing loading and flight muscle mass are only shown for males as females are flightless.

**Figure S3:**
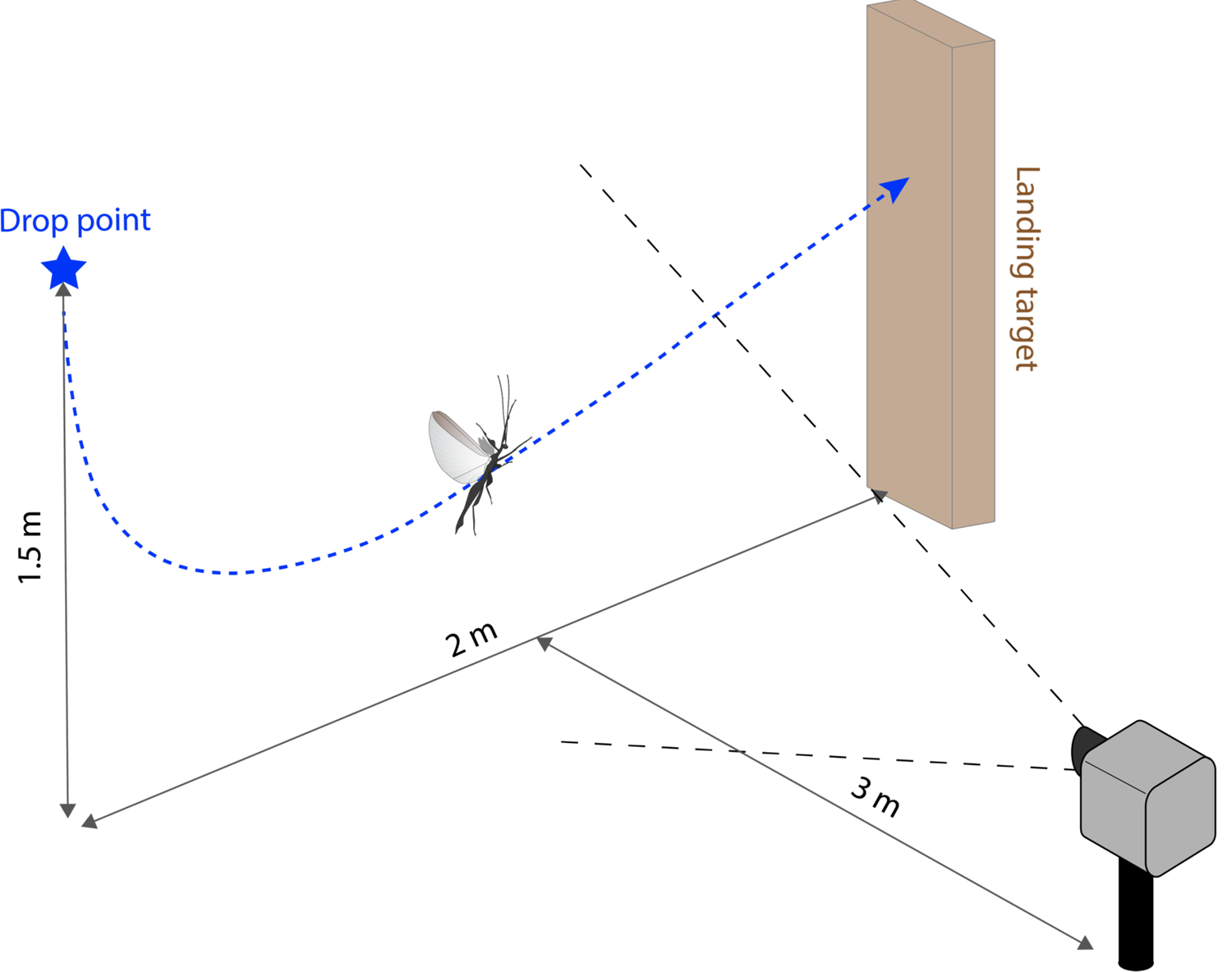
Set-up for flight trials.

**Figure S4:**
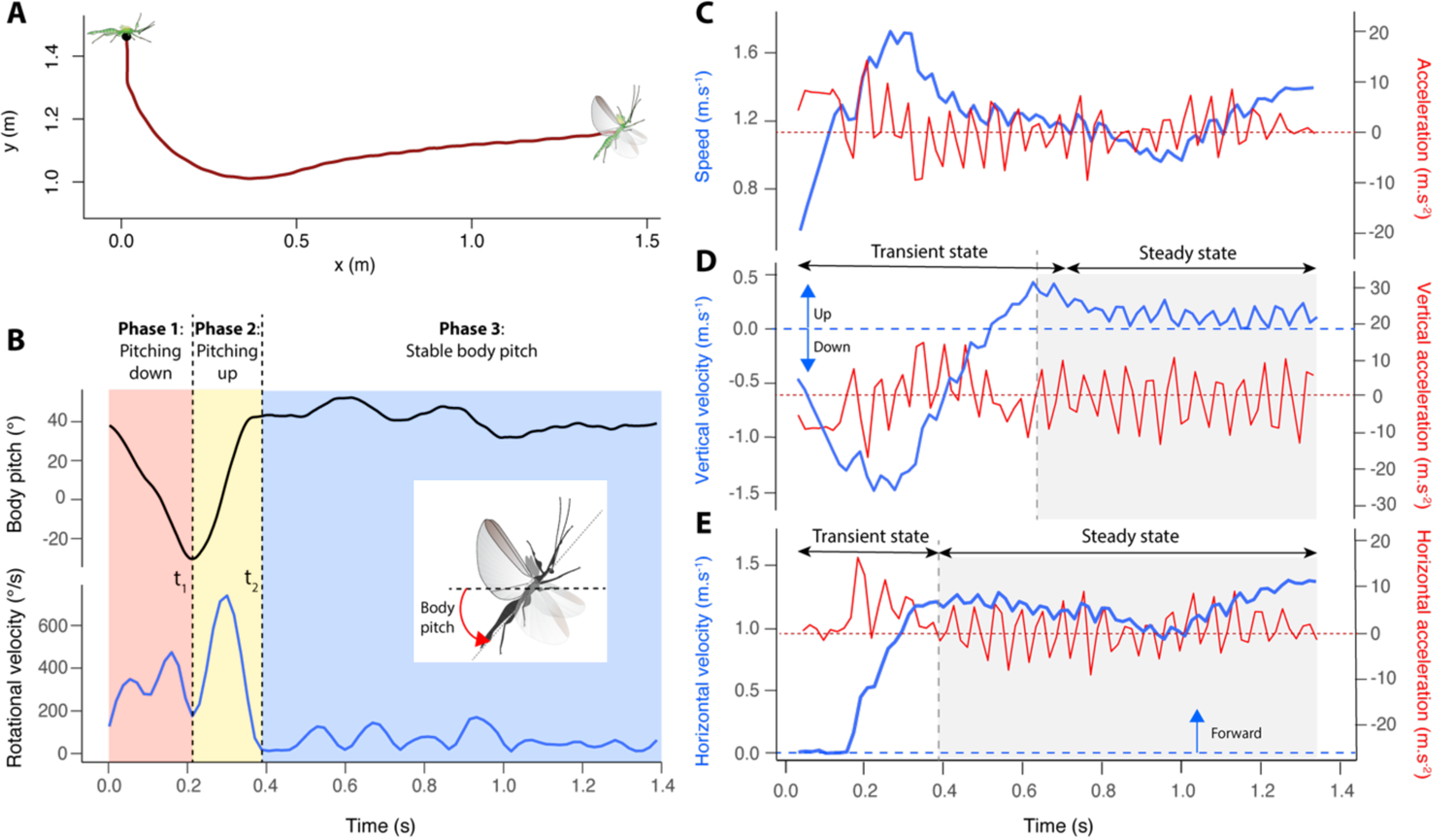
Analysis of a male *P. philippinicum* flight trial. **A**: Smoothed trajectory. Cartoons show the typical postures of the insect at the beginning and end of the trial. **B**: Body pitch and body pitch rotational velocity as a function of time. Colors indicate different phases during the trial defined by t_1_ – i.e., to the time when body pitch reaches its absolute minimum – and t_2_ – i.e., the time when body pitch stabilizes. **C**: Body speed and overall acceleration as a function of time. **D**: Vertical body velocity and acceleration as a function of time. Positive values indicate an upward velocity or acceleration. **E**: Horizontal body velocity and acceleration as a function of time. Positive values indicate a forward velocity or acceleration. In **D** and **E**, the boundary between the transient and steady state corresponds to the time when velocity (vertical or horizontal) stabilizes.

**Figure S5:**
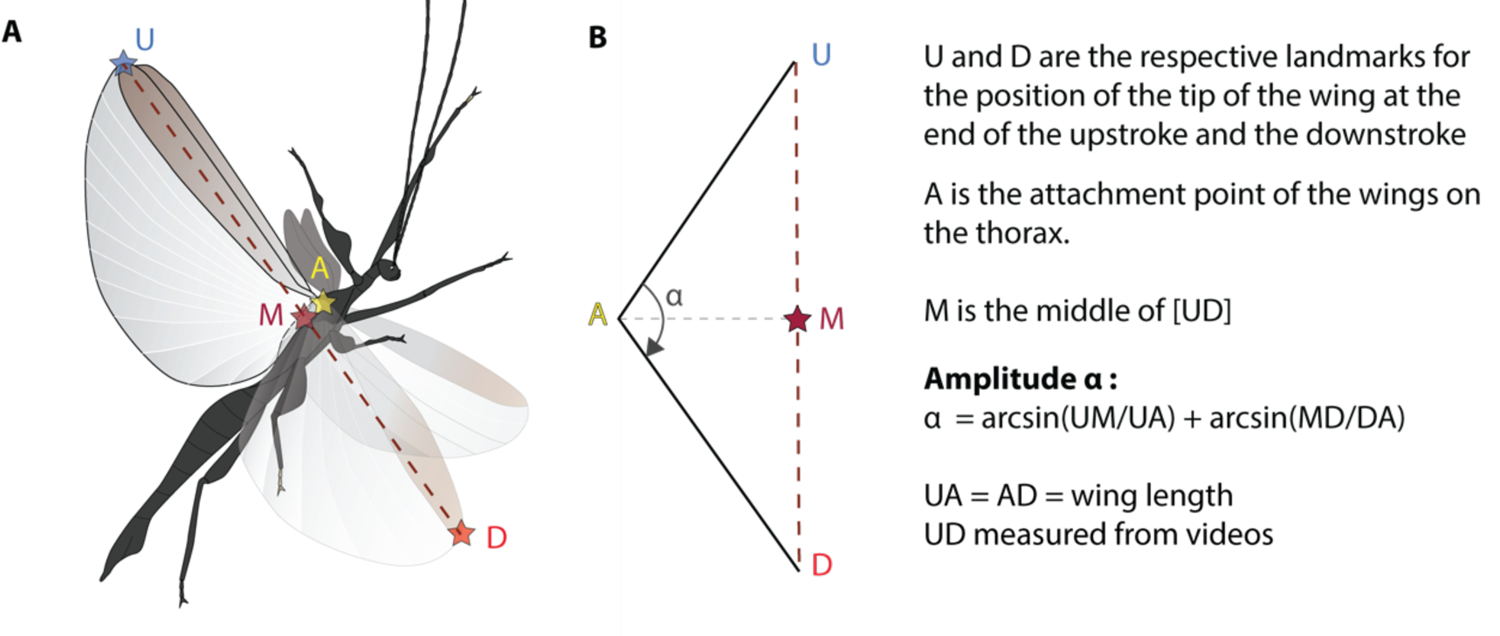
Calculation of wing stroke amplitude. **A:** side view of the animal with the relevant landmarks. **B:** landmarks in frontal view.

**Figure S6:**
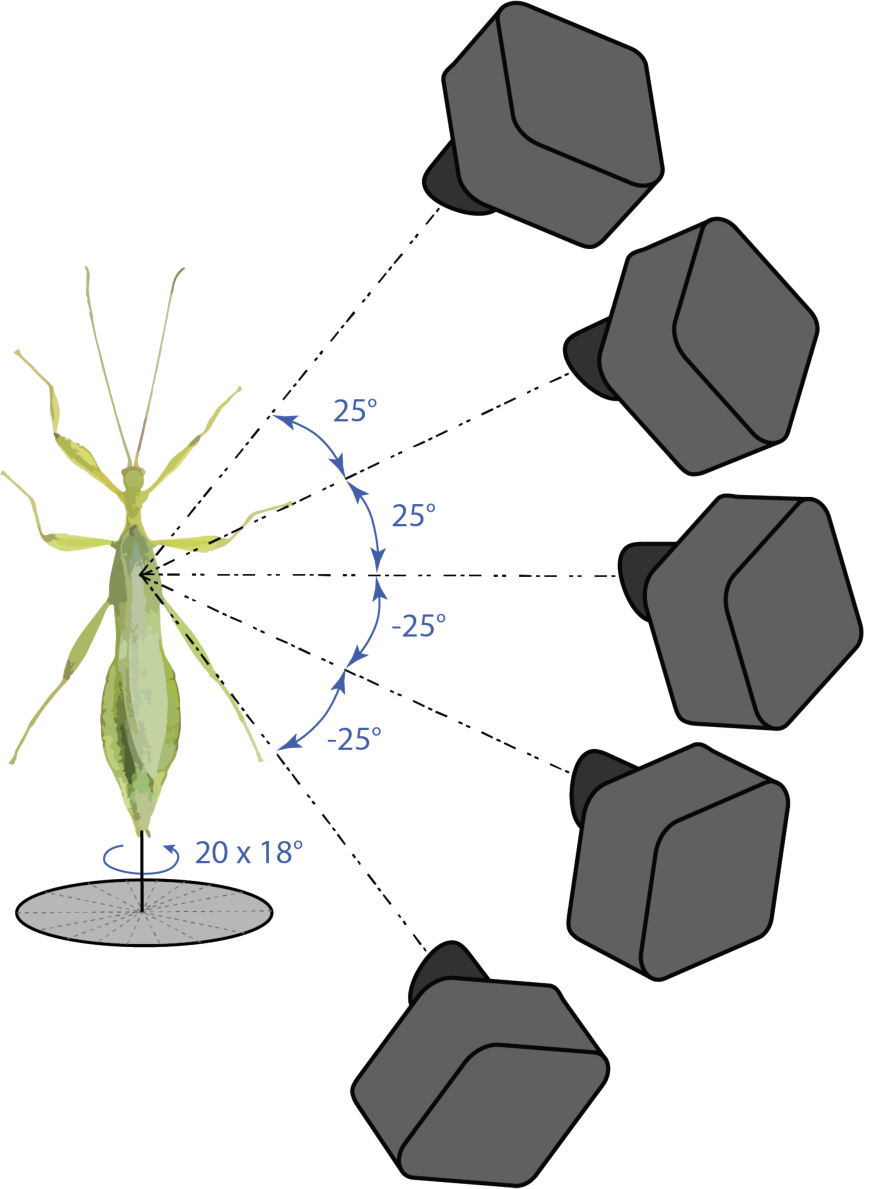
Acquisition of multiple 2D photographs from different angles of a male mounted on a pin to reconstruct a single 3D model using photogrammetry.

**Figure S7:**
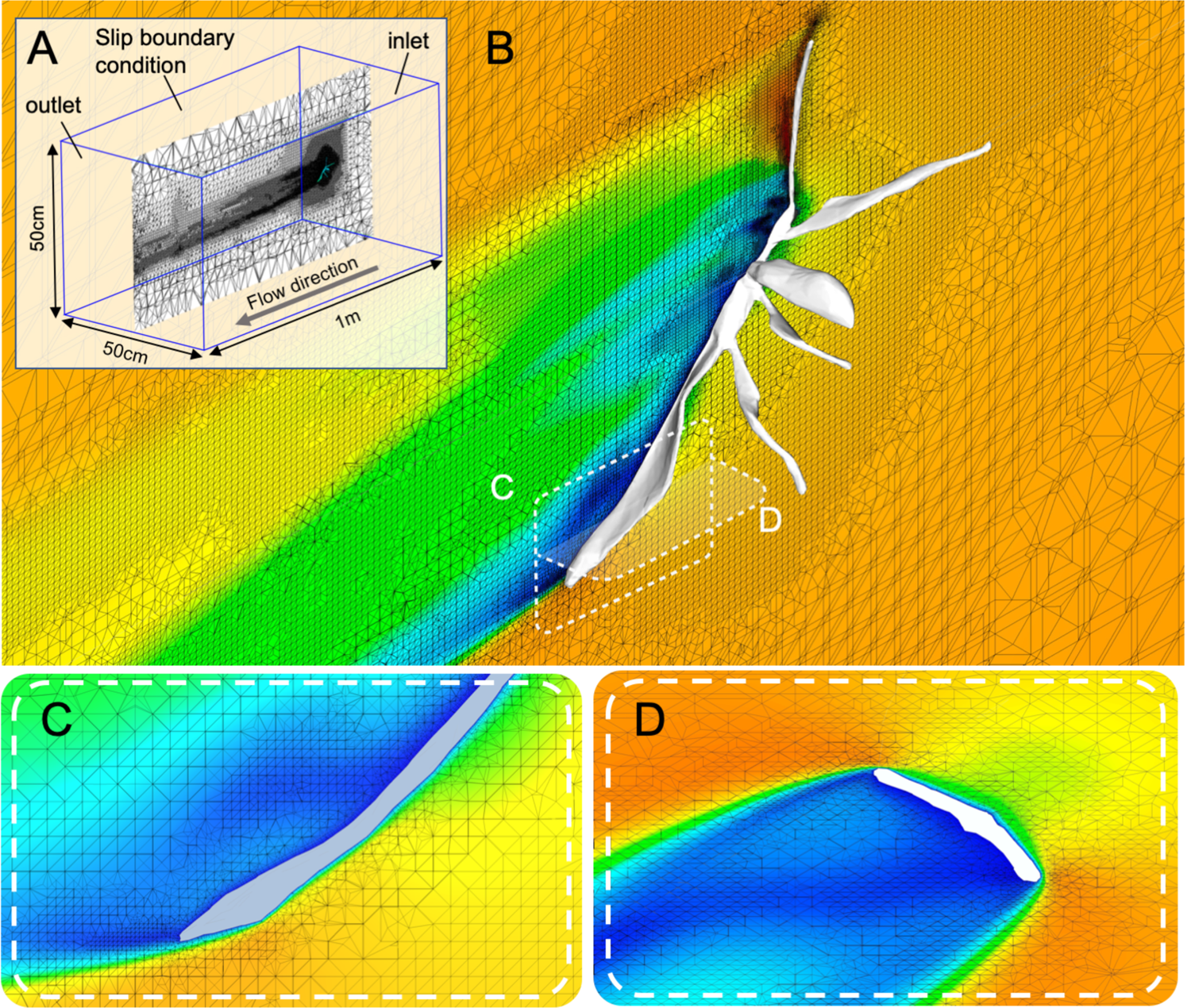
Leaf insect model and CFD simulation results. **A:** Meshed CFD domain. **B:** Detail of the mesh around the phasmid model along the median sagittal plane. **C:** Detail of the mesh around the tip of the abdomen along the median sagittal plane. **D:** Detail of the mesh around the middle of the abdomen along a cross-sectional plane. Colors in **B-D** represent air velocity (see scale in figure 6).

**Video S1:** Example of a flight trajectory from a relatively light male (374.0 mg)

**Video S2:** Example of a flight trajectory from a relatively heavy male (504.7mg)

**Video S3:** Detail of a male leaf insect ascending.

